# Membrane surfaces regulate assembly of a ribonucleoprotein condensate

**DOI:** 10.1101/2021.04.24.441251

**Authors:** Wilton T. Snead, Therese M. Gerbich, Ian Seim, Zhongxiu Hu, Amy S. Gladfelter

**Author notes:** To whom correspondence should be addressed: Amy S. Gladfelter.

## Abstract

Biomolecular condensates organize biochemistry in time and space, yet little is known about how cells control either the position or scale of these assemblies. In cells, condensates often appear as dispersed, relatively small assemblies that do not grow (coarsen) into a single droplet despite their propensity to coalesce. Here we report that ribonucleoprotein condensates of the Q-rich protein Whi3 interact with the endoplasmic reticulum, prompting us to hypothesize that membrane association controls the position and size of condensates. Reconstitution of Whi3 condensates on supported lipid bilayers reveals that association with a diffusive lipid surface promotes condensation at both physiological ionic strength and protein concentration. Notably, these assemblies rapidly arrest, matching size distributions seen in cells. The timing of the arrest is influenced by the ordering of protein-protein and protein-RNA interactions and controlled by the slow diffusion of complexes induced by the membrane. This slowed diffusion limits both transfer of small protein-RNA complexes between condensates and their coalescence, thus driving coarsening to arrest. Our experiments reveal a tradeoff between locally-enhanced protein concentration at membranes, which favors condensation, and an accompanying reduction in diffusion, which restricts coarsening. Thus, membranes can maintain a population of small condensates in the absence of active mechanisms. Given that many condensates are bound to endomembranes, we predict that the biophysical properties of lipid bilayers are key for controlling condensate sizes throughout the cell.

**One sentence summary:** Assembly on a membrane surface positions and scales biomolecular condensates by controlling relative diffusion rates of proteins and nucleic acids.

## Introduction

Compartmentalization of biochemistry is critical for diverse cell functions. Biomolecular condensates, composed of a concentrated assembly of protein and nucleic acid, are emerging as centers of compartmentalization throughout the cell (*1*). Many biomolecular condensates form via the process of liquid-liquid phase separation (LLPS), in which transient, multivalent interactions among proteins and nucleic acid polymers facilitate the formation of a condensed phase. LLPS occurs when the bulk concentration exceeds a saturation concentration, above which molecules spontaneously partition into dense and dilute phases (*2, 3*). Condensates formed by LLPS can display properties of liquid-like droplets and dynamically exchange molecules with the surrounding environment (*4*).

LLPS is characterized by distinct regimes of condensate growth and coarsening (*2*). In the growth regime, the dilute phase becomes depleted as condensates approach the equilibrium dense phase concentration. In the coarsening regime, condensates mature and increase in size through processes such as Ostwald ripening and Brownian motion coalescence (*2, 5*), both of which are characterized by power law dynamic scaling over time with an exponent of 1/3 (*2, 6-8*). Equilibrium thermodynamics predicts that coarsening will eventually result in a single large droplet that minimizes the system’s interfacial energy as opposed to a distribution of smaller droplets with higher interfacial energy. Indeed, protein and RNA condensates frequently coarsen into large, micrometer-size droplets when reconstituted *in vitro* (*5, 9-12*). However, cells are not in equilibrium, and condensates within living cells are often an emulsion of diffraction-limited puncta or small droplets that do not increase beyond a certain size (*12-16*). The cellular mechanisms that restrict coarsening and control condensate size remain poorly understood.

Recent studies have shown that diverse biomolecular condensates are associated with and potentially regulated by membrane surfaces (*17-19*). For example, the plasma membrane controls formation of condensates involved in immune cell activation (*20-23*), actin assembly (*24, 25*), and presynaptic active zones (*26, 27*). Many condensates also require associations with endomembrane surfaces for proper transport and subcellular positioning (*28-30*), suggesting that interactions with membrane-bound organelles are important for condensate formation. For example, phase separation of autophagy-related proteins at the vacuole regulates assembly of the autophagosome (*31*), and condensate wetting on autophagosomal membranes is key for condensate removal (*32*). While the interactions between condensates and membrane-bound organelles are still being uncovered, growing evidence indicates that the assembly and morphology of many condensates are closely linked with the endoplasmic reticulum (ER) (*16, 18, 33-35*).

Our group discovered that the RNA-binding, glutamine (Q)-rich protein Whi3 forms spatially-distinct condensates that serve essential physiological functions in the multinucleate fungus *Ashbya gossypii* (*12, 36-38*). Specifically, Whi3 forms punctate, diffraction-limited condensates containing either cyclin or formin RNA transcripts, which are positioned near nuclei and sites of polarized growth, respectively. These condensates are essential regulators of the cell cycle and cell polarity. *In vitro*, Whi3 and its binding RNAs form condensates that coarsen into large, macroscopic droplets (*12, 38*). However, it is unknown how *Ashbya* cells restrict Whi3 condensate coarsening and maintain a population of small puncta in distinct cellular locations.

In this work, we find that Whi3 condensates stably associate with the ER, suggesting that endomembrane surfaces may contribute to regulating Whi3 condensate assembly. In support of this hypothesis, we find that Whi3 recruitment to synthetic membranes substantially reduces the bulk protein concentrations required for condensation compared to solution droplets. However, membranes also rapidly suppress condensate coarsening by reducing protein diffusion compared to solution. Surprisingly, we find that heterotypic Whi3-RNA interactions reduce diffusion even further by trapping proteins within slow-diffusing RNA clusters. As a result, coarsening of membrane-associated Whi3 condensates arrests even faster in the presence of RNA. Moreover, recruitment of Whi3 to membrane-tethered RNA, which enables protein-RNA interactions to occur earlier during assembly, drives the formation of punctate assemblies which do not coarsen into micrometer-scale droplets. The size distribution of these puncta is similar to that of native Whi3 assemblies in *Ashbya* cells. Our results reveal that the biophysical properties of membranes can both promote protein-RNA condensate assembly and control condensate size, suggesting that membrane attachment is likely a key mechanism for controlling the formation of diverse condensates throughout the cell.

## Results

### Whi3 condensates persistently associate with the endoplasmic reticulum

*Ashbya gossypii* Whi3 is a 79 kDa protein containing a low-complexity, Q-rich region followed by a structured RNA recognition motif. Our previous work found that Whi3 undergoes a phase transition to form liquid-like droplets in the presence of its binding RNAs (*12, 38*). This phase transition is mediated by interactions among the Q-rich region as well as RNA binding. Specifically, combining Whi3 with target RNAs *in vitro* yields micrometer-sized liquid-like droplets (*12, 38*). However, Whi3-based assemblies in cells appear as small puncta rather than micrometer-sized droplets. We set out to understand this scaling difference between *in vivo* and *in vitro* Whi3 condensates, and to identify mechanisms that control droplet coarsening in cells.

Some target RNAs of Whi3 are enriched in the vicinity of nuclei and at the cell cortex (*36, 37*), prompting us to hypothesize that that Whi3 condensates may be associated with the endoplasmic reticulum (ER), which is associated with both nuclei and the cortex. To test this, we created an *Ashbya* strain in which Whi3 was tagged endogenously with tdTomato (*39*) and a GFP-tagged ER marker, the translocon component Sec63, was expressed from a plasmid. Whi3 puncta in these cells appeared co-localized with diverse ER structures, including tubules and nuclear-associated ER (Fig. 1A). To quantify co-localization, we detected Whi3 puncta (*40*) and determined the associated, local fluorescence intensity in the ER channel at each detected position (see methods and Fig. S1). This analysis revealed that approximately 81% of Whi3 puncta were co-localized with the ER, compared to approximately 50% of puncta appearing co-localized with the ER by chance when simulated as a random distribution (Fig. 1B). When compared to 50 different random distributions, we found that observed Whi3 puncta consistently showed a greater local intensity in the ER channel compared to random distributions, by a factor of 1.62 ± 0.36 s.d. (Fig. 1C). Together, these data indicate that Whi3 assemblies are consistently more co-localized with the ER than expected by chance.

**Fig. 1.**
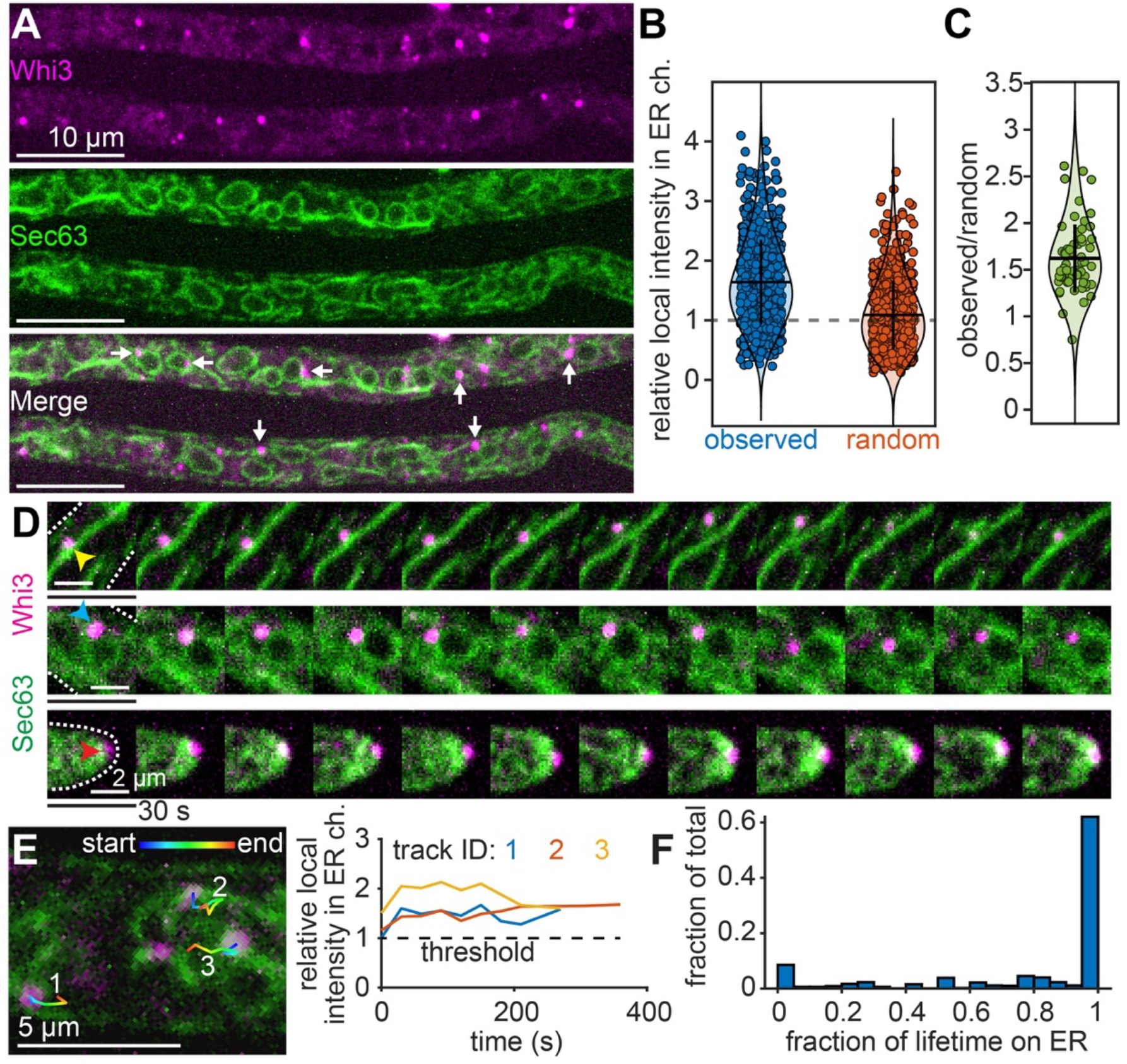
Whi3 puncta persistently associate with the endoplasmic reticulum in *Ashbya gossypii*. (**A**) *Ashbya* hyphae expressing Whi3-tdTomato and Sec63-GFP (ER marker). Example Whi3 puncta showing ER co-localization are indicated by white arrows. (**B**) Local intensity in the ER channel at detected Whi3 puncta, expressed as a fraction of the median intensity of the ER channel throughout the hypha, after masking and background subtraction (see methods). Blue points: observed Whi3 positions, orange points: randomized Whi3 positions. n = 642 puncta. Dashed line indicates the threshold for co-localization, corresponding to the median intensity of the ER channel. (**C**) Ratio of the average local intensity in the ER channel at detected Whi3 puncta within a hypha relative to randomized Whi3 positions. Each data point represents the average of 50 random distributions per hypha, n = 60 hyphae (see methods). Horizontal bars in (B) and (C) represent mean, vertical bars represent standard deviation. (**D**) Montages of Whi3 puncta associated with ER tubule (top, yellow arrowhead), nuclear-associated ER (middle, blue arrowhead), and hyphal tip (bottom, red arrowhead). White dashed lines in first frame indicate cell periphery. Montages span 6 min, 30 s/frame. (**E**) First frame from a time-lapse showing ER (green) and Whi3 (magenta) with three overlaid Whi3 tracks. Plot shows the associated, local intensity in the ER channel at each time point of the three tracks, expressed as a fraction of the median intensity of the ER channel throughout the hypha. All three tracks spend the entire lifetime co-localized with the ER. (**F**) Histogram of Whi3 tracks, binned according to the fraction of track lifetime co-localized with the ER. n = 769 tracks.

To assess whether Whi3 puncta associate with the ER transiently, or if puncta remain stably tethered, we visualized and tracked Whi3-ER co-localization over time (see methods and Fig. S2). We observed that the movement of Whi3 puncta frequently tracked with the movement of the ER (Fig. 1D and Movie S1) and that the majority of Whi3 tracks were strongly associated with the ER for most of the track lifetime (Fig. 1F). In particular, 74% of tracks spent 75% or more of the lifetime co-localized with the ER, while 62% of tracks spent the entire lifetime co-localized with the ER (Fig. 1F). Collectively, these findings indicate that a substantial fraction of Whi3 puncta are stably tethered to the ER over time, consistent with a previous report that protein-RNA condensates associate with the ER in mammalian cells (*16*).

These observations prompted us to hypothesize that the ER surface may contribute to Whi3 condensate assembly. Specifically, we imagined that membranes may lower the concentration of protein required for phase separation (*9, 24*) and/or supply factors that limit coarsening. To examine the biophysical properties of membranes that contribute to condensate formation and size distributions seen in cells, we reconstituted Whi3 condensate formation on synthetic membrane surfaces.

### Whi3 rapidly forms condensates on synthetic membranes

To analyze how association with membranes may control condensate assembly, we reconstituted Whi3 condensates on planar, supported lipid bilayers (SLBs). Because Whi3 does not contain an obvious lipid interaction domain, Whi3 was recruited to SLBs via a hexa-histidine (6his)-nickel nitrilotriacetic acid (NTA-Ni) interaction. We first asked whether membrane surfaces may promote condensate assembly by increasing the local concentration of Whi3 relative to the solution. We therefore added Whi3 protein to SLBs at a concentration of 50 nM, comparable to previous measurements of the soluble Whi3 concentration in *Ashbya* cells, but several orders of magnitude lower than concentrations required to drive protein-only phase separation *in vitro* (*12*). SLBs were imaged using TIRF microscopy, as portrayed in Fig. 2A. We found that circular, patch-like Whi3 condensates formed on SLBs within approximately 2 min of protein addition (Fig. 2B; Movie S2). Notably, these experiments were performed in a buffer with physiological ionic strength (150 mM KCl), while previous experiments required a lower ionic strength (75 mM) and micromolar concentrations of Whi3 to observe protein-only demixing in solution (*12*). Therefore, association with a membrane surface appears to dramatically shift the phase boundary, reducing the bulk Whi3 concentration and salt barrier required for condensate assembly in the absence of RNA.

**Fig. 2.**
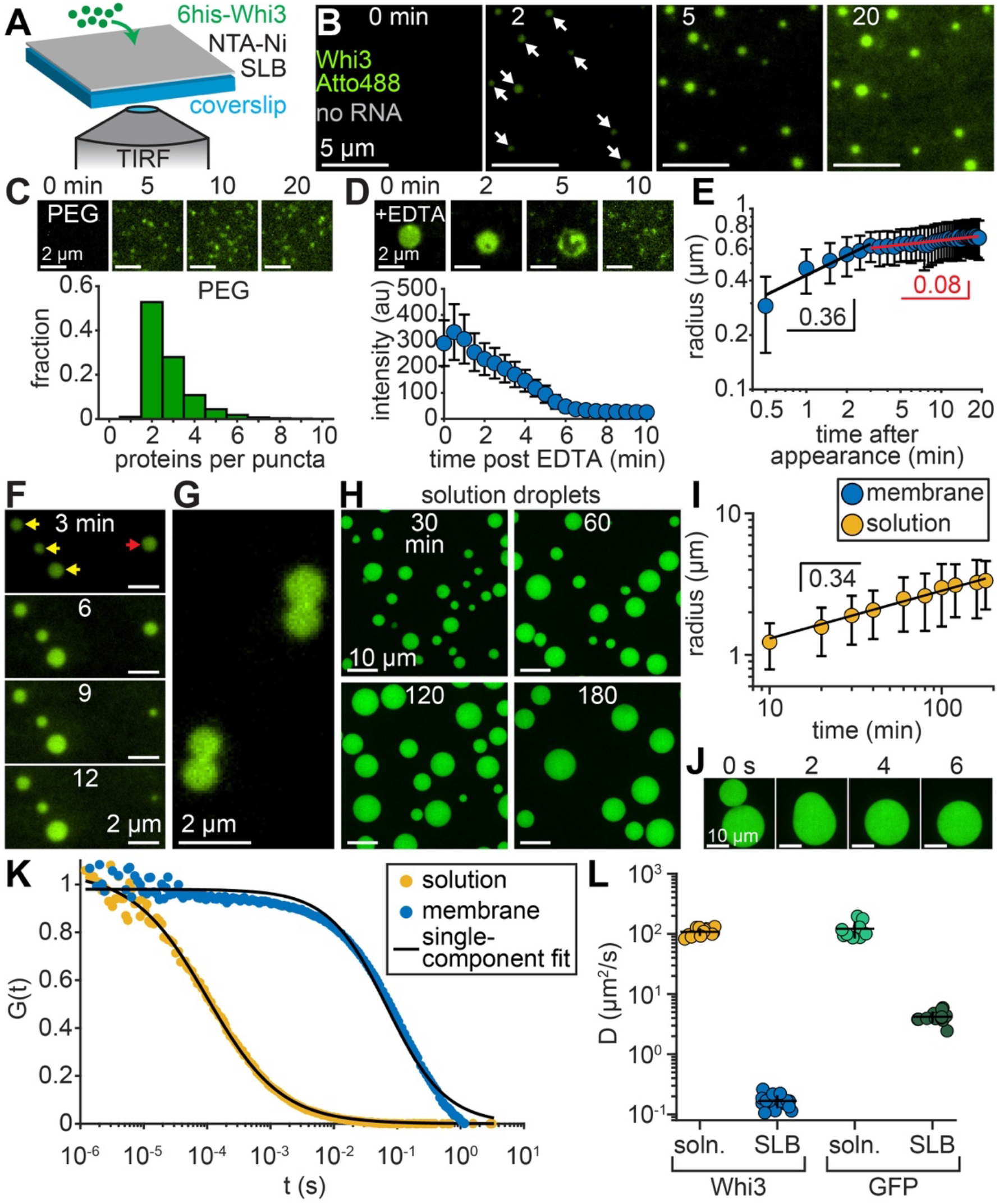
Membrane-associated Whi3 condensates rapidly assemble and arrest. (**A**) Schematic of SLB experiment. (**B**) Time-lapse of SLB after addition of 50 nM Whi3-Atto488. White arrows indicate condensates apparent after 2 min. (**C**) Top: Time-lapse of PEG surface after addition of 50 nM Whi3. Bottom: Distribution of the number of Whi3 proteins per puncta on PEG surfaces. (**D**) Top: Time-lapse of membrane-associated condensate, formed with 50 nM Whi3, after addition of 10 mM EDTA. Bottom: Average condensate intensity as a function of time after addition of 10 mM EDTA. (**E**) Average condensate radius as a function of time after condensates first appear, formed with 50 nM Whi3. Black and red lines show fitted power law functions to early and late coarsening regimes, respectively, with indicated scaling exponents. (**F**) Time-lapse of SLB at the indicated times after addition of 1 µM Whi3. Condensates that grow or shrink in size over time are indicated by yellow or red arrows, respectively. (**G**) Condensates on SLB, approximately 20 min after addition of 50 nM Whi3, showing no fusion or rounding upon contact. (**H**) Solution droplets formed with 41 µM Whi3 at the indicated times after assembly, induced by lowering the KCl concentration to 75 mM. Images are maximum intensity projections from confocal z-stacks. All images contrasted equally. (**I**) Average radius of solution droplets as a function of time after assembly, formed with 41 µM Whi3. Black line shows fitted power law function with indicated scaling exponent. (**J**) Time-lapse of rapid droplet fusion, 4 h after assembly. (**K**) Average, normalized FCS traces of Whi3 diffusion in solution or at membrane. Black curves indicate fits to single-component diffusion model. (**L**) Diffusion coefficients of Whi3 and GFP in solution or at membranes, estimated from fitting FCS traces to a single-component diffusion model. n > 10,000 puncta in (C); n = 10 condensates in (D); n > 50 tracked condensates in (E); n > 400 droplets per data point in (I); n > 9 FCS traces per condition in (K-L). All data pooled from at least three independent experiments. All error bars represent standard deviation. SLB membrane composition: 96 mol% DOPC, 4 mol% DGS NTA-Ni.

We next tested if this shift in the phase boundary was simply driven by an interaction with a surface, or if the diffusive properties of a membrane were required. We found that condensates did not appear when 50 nM Whi3 was added to an immobile, PEG-coated surface (Fig. 2C). Rather, proteins remained primarily as dimers or trimers on PEG surfaces, as revealed by particle detection and single molecule calibration (Fig. 2C and S3). Thus, small Whi3 oligomers form in solution and on passivated surfaces at this concentration, but higher-order assemblies or condensates do not. Moreover, addition of 10 mM EDTA to SLBs, which chelates Ni^2+^ ions and disrupts Whi3-membrane binding, induced condensate disassembly within approximately 6 min (Fig. 2D). Thus, recruitment to a diffusive lipid surface was required for Whi3 condensate formation.

### Coarsening is rapidly suppressed at membrane surfaces

In addition to promoting assembly in a distinct phase regime compared to solution condensates (*12*), membrane-associated condensates also appeared to plateau in size within approximately 5 min of protein addition (Fig. 2 B and E; Movie S2). Specifically, the average condensate radius over time followed power law scaling for approximately 3 min after condensates first appeared, with a dynamic scaling exponent of 0.36 ± 0.12 (95% CI of fit) (Fig. 2E). This early coarsening period was followed by a second, longer regime with a substantially slower power law scaling exponent of 0.08 ± 0.01 (95% CI of fit) (Fig. 2E). The slow scaling during this second regime indicates that condensate size was almost completely arrested.

The early scaling regime in Fig. 2E is consistent with coarsening mechanisms such as diffusion-limited Ostwald ripening or Brownian motion coalescence, which predict condensate size scaling over time with a power law exponent of 1/3 (*2, 5-8*). We speculate that Ostwald ripening may have contributed to coarsening during this early period, as some condensates occasionally dissolved while others grew larger (Fig. 2F). By contrast, condensates did not fuse and round upon contact (Fig. 2G), indicating that Brownian motion coalescence did not contribute to coarsening. This finding also suggests that condensates may have rapidly acquired viscoelastic properties that prevented fusion and may have contributed to the suppression of coarsening (*41*) (Fig. 2G). Collectively, these experiments reveal that even though membrane surfaces promote rapid assembly of protein-only Whi3 condensates, coarsening is also quickly suppressed.

### Solution droplets coarsen for substantially longer compared to membrane-associated condensates

To test the hypothesis that membrane surfaces hinder coarsening, we compared the coarsening dynamics of Whi3 condensates formed in solution and at membrane surfaces. To assemble Whi3 droplets in solution, the buffer ionic strength was reduced (75 mM KCl) in the presence of 41 µM Whi3, which promoted Whi3 protein-only phase separation (*12*). We found that solution droplets coarsened continually for at least 3 h (Fig. 2H), and the average droplet radius increased monotonically from 1.23 ± 0.44 to 3.35 ± 1.26 µm s.d. between 10 and 180 min after droplet assembly (Fig. 2I). The coarsening dynamics during this period followed a clear power law trend with a scaling exponent of 0.34 ± 0.04, 95% CI of fit, consistent with the power law dynamic scaling exponent of 1/3 expected for liquid-like droplet coarsening (Fig. 2I) (*2, 5-8*). Brownian motion coalescence likely contributed to coarsening here, as droplets rapidly fused and rounded up even 4 h after assembly (Fig. 2J). By contrast, membrane-associated Whi3 condensates only appeared to coarsen for approximately 3 min before arresting (Fig. 2E), and the average condensate radius remained relatively constant when the bulk Whi3 concentration ranged from 20-500 nM (Fig. S4 A-B). Collectively, these data suggest that membrane surfaces dramatically accelerate an arrest in coarsening compared to solution droplets, thereby stabilizing condensates within a relatively narrow size range over a broad range of concentrations.

### Membrane surfaces reduce Whi3 diffusion

How do membrane surfaces hinder coarsening? Given that the bulk Whi3 concentration in these experiments was 50 nM, well below the saturation concentration for phase separation in solution, we predicted that the predominant protein source for condensate growth came from the membrane. Specifically, the membrane-bound protein density at the approximate time of condensate appearance was 217 proteins/µm^2^, equivalent to a local concentration of approximately 80 µM (Fig. S4C; see methods). We hypothesized that despite this high local concentration, membrane-bound protein may have been less diffusive in comparison to free-diffusing protein in solution (*42, 43*), which could inhibit protein incorporation into condensates.

To test the idea that differential diffusion rates can control coarsening, we used fluorescence correlation spectroscopy (FCS) to examine the diffusion of Whi3 in solution or bound to SLBs (*43*). We found that membrane-bound Whi3 diffused dramatically slower compared to Whi3 in solution (Fig. 2K), with a more than 600-fold reduction in the average diffusion coefficient from 108 ± 16 to 0.17 ± 0.04 µm^2^ s^-1^ s.d. in solution and at membranes, respectively (Fig. 2L). By comparison, the diffusion coefficient of GFP, which does not self-assemble like Whi3, was reduced by approximately 30-fold when membrane-bound (Fig. 2L and S5). Therefore, the diffusion of membrane-bound Whi3 was reduced more than would be expected for proteins that do not self-assemble. Specifically, Whi3 recruitment to membranes likely promoted the assembly of higher-order oligomers that diffused slower compared to monomers. Indeed, FCS traces of membrane-bound Whi3 deviated from a single-component diffusion model (Fig. 2K), indicating that a mixture of Whi3 oligomers of varying size and diffusion likely contributed to the spread in our measurements.

Our experiments to this point show that direct recruitment of Whi3 to membrane surfaces can dramatically shift the phase boundary for Whi3 protein-only condensate assembly. Once formed, these membrane-associated condensates rapidly arrest, potentially due to a reduction in protein diffusion. However, Whi3 condensates assemble with RNA in *Ashbya* cells (*36-38*), and RNA promotes Whi3 phase separation *in vitro* (*12, 38*). We next asked if, and how, RNA may influence the coarsening dynamics and size distributions of Whi3 condensates at membranes.

### RNA suppresses coarsening at membranes

To begin analyzing the role of RNA in the assembly of membrane-associated condensates, a Whi3-interacting RNA, the cyclin *CLN3*, was added in reactions prior to addition of Whi3 protein. *CLN3* was included at a concentration of 100 pM, comparable to a previous estimate of the *CLN3* RNA transcript concentration in *Ashbya* cells (*36*). Similar to RNA-free experiments (Fig. 2B), circular, patch-like Whi3 condensates formed within 2 min of protein addition (Fig. 3A), again reaching a plateau within approximately 5 min (Fig. 3B; Movie S3). Throughout the course of assembly, RNA was progressively recruited from solution, appearing as puncta distributed around the membrane surface (Fig. 3A; Movie S3). However, RNA was clearly excluded from the interior of membrane-associated condensates, instead interacting primarily with the periphery (Fig. 3 A and D; Movie S4). Condensates also frequently nucleated with no RNA association (Fig. S6A), suggesting that initial assembly was driven primarily by Whi3 homotypic interactions, while interactions with RNA were restricted to condensate edges. Indeed, the early dynamic scaling exponents prior to condensate arrest were 0.35 ± 0.06 and 0.36 ± 0.12 (95% CI of fits) in the presence and absence of RNA, respectively (Fig. S6B), indicating that the initial assembly and coarsening dynamics were independent of RNA.

**Fig. 3.**
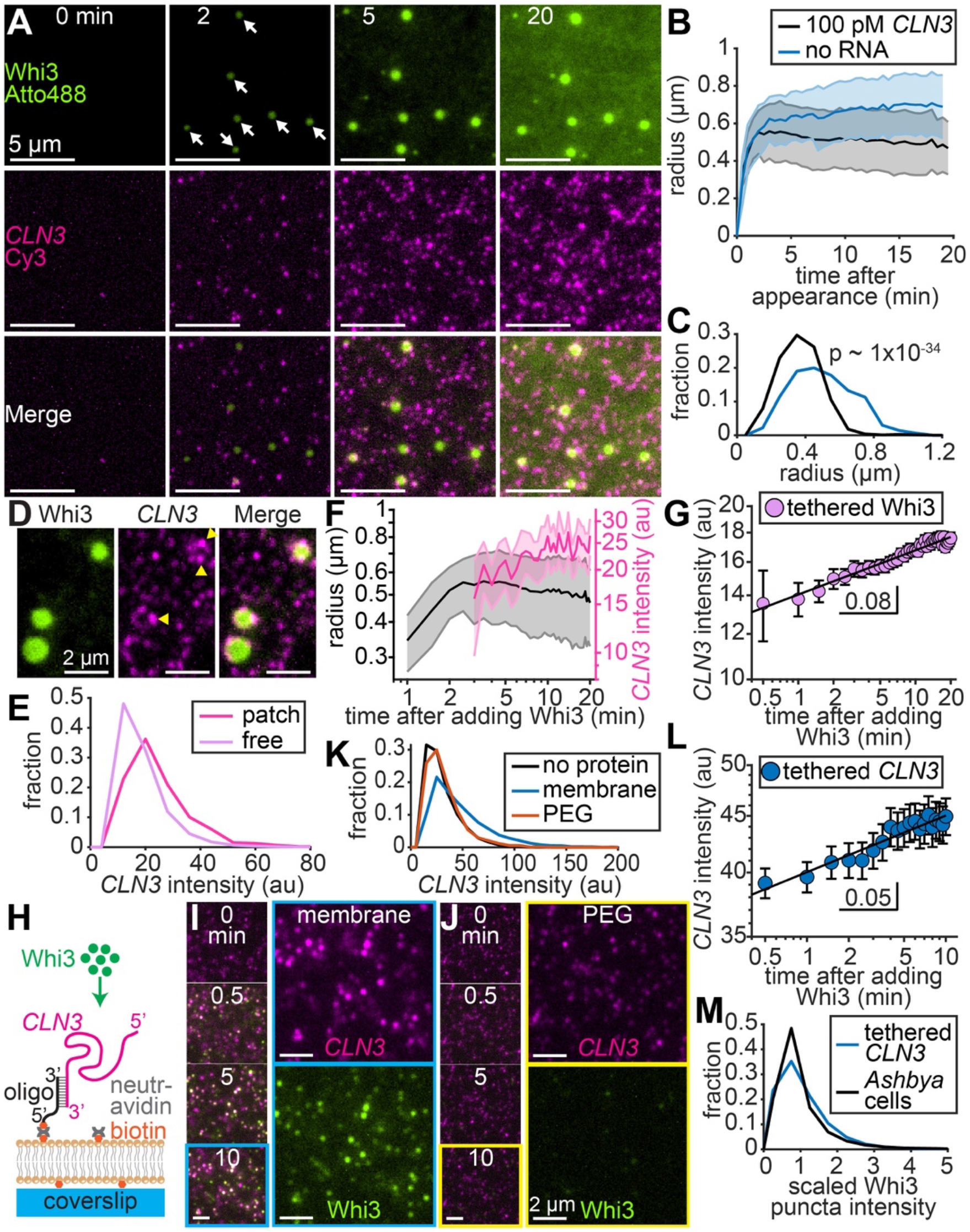
RNA suppresses condensate coarsening at membranes by forming small clusters. **(A)** Time-lapse of SLB with *CLN3*-Cy3 in solution prior to Whi3 addition. Frames show time after addition of 50 nM Whi3-Atto488. Final *CLN3* concentration is 100 pM. White arrows indicate condensates apparent after 2 min. (**B**) Average condensate radius as a function of time after condensates first appear, formed with 50 nM Whi3 in the presence and absence of RNA. Blue curve repeated from Fig. 2E. (**C**) Distributions of condensate radii formed with 50 nM Whi3, in the presence and absence of RNA, 20-30 min after condensate assembly. Indicated p-value from 2-sample Kolmogorov-Smirnov test. (**D**) *CLN3* RNA excluded from interior of membrane-associated condensates formed with 50 nM Whi3 and 100 pM *CLN3*. Yellow arrowheads indicate bright *CLN3* puncta at condensate edges. (**E**) Intensity distributions of *CLN3* puncta associated with Whi3 condensate edges (“patch”) or on the surrounding membrane (“free”), 20-30 min after addition of 50 nM Whi3. (**F**) Overlay of average condensate radius, repeated from (B) and average intensity of *CLN3* puncta at condensate edges as a function of time after addition of 50 nM Whi3. (**G**) Average intensity of “free” *CLN3* puncta recruited by membrane-bound Whi3 as a function of time after addition of 50 nM Whi3. Black line shows fit to a power law function with indicated exponent. (**H**) Schematic of membrane-tethered RNA experiment. (**I-J**) SLB (I) and immobile, PEG-coated surface (J) with tethered *CLN3* at indicated times after addition of 50 nM Whi3. Time lapse images show merged *CLN3* and Whi3 channels, while the cyan (I) or yellow (J) boxed frames show separate channels at 10 min time point. *CLN3* and Whi3 channels contrasted equally. (**K**) Intensity distributions of membrane or PEG surface-tethered *CLN3* puncta before and after addition of 50 nM Whi3. (**L**) Average intensity of membrane-tethered *CLN3* puncta as a function of time after addition of 50 nM Whi3. Black line shows fit to a power law function with indicated exponent. (**M**) Distributions of scaled Whi3 puncta intensity from *Ashbya* cells and membrane-tethered *CLN3* experiments, after clustering by 400 nM Whi3. Scaled distributions obtained by dividing by the distribution means. n > 50 tracked condensates per condition in (B); n > 500 condensates per condition in (C); n = 467 and 24,954 patch-associated and free RNA puncta in (E); n ranges from 7 - 48 puncta per data point in (F – magenta curve); n ranges from 44 - 2,264 puncta per data point in (G); n > 3,800 puncta per condition in (K); n > 1,300 puncta per data point in (L); n = 18,503 and 867 puncta in tethered *CLN3* experiment and live *Ashbya* cell distributions, respectively, in (M). All data pooled from at least three independent experiments. Error bars in (B) and (F – black curve): standard deviation; (F – magenta curve), (G), and (L): 95% CI. SLB membrane composition in (A-G): 96 mol% DOPC, 4 mol% DGS NTA-Ni; in (I, K-M): 99 mol% DOPC, 1 mol% DOPE cap-biotin.

Although RNA did not influence the initial assembly dynamics of membrane-associated Whi3 condensates, we found that condensates arrested faster in the presence of RNA (Fig. 3B) and the distribution of condensate radii after 20 min of assembly was significantly smaller on average and more monodispersed compared to RNA-free condensates (Fig. 3C). These findings suggest that recruitment of RNA from solution can accelerate the arrest of protein-only condensate coarsening. How might RNA suppress coarsening?

### RNA recruited by membrane-bound Whi3 forms clusters that resist coarsening

To examine how interactions with *CLN3* RNA impact coarsening, we analyzed the dynamics of RNA recruitment by membrane-bound Whi3. We found that the intensity of RNA puncta across the membrane increased with time (Fig. 3A; Movie S3), suggesting that Whi3 clustered RNA into higher-order assemblies. Notably, many RNA puncta associated with the periphery of condensed Whi3 patches were brighter compared to “free” puncta on the surrounding membrane (Fig. 3 D and E; Movie S4). These RNA puncta began to appear and cluster into brighter assemblies at patch edges at roughly the same time that patch coarsening arrested (Fig. 3F and S6 C-D), suggesting that edge-associated RNA clusters may have contributed to the suppression of coarsening. Interestingly, recent work found that an accumulation of negative charge at condensate interfaces can reduce droplet surface tension (*44*), a key determinant of the coarsening rate. It is possible that *CLN3* clusters at condensate edges may have suppressed coarsening by a similar mechanism.

Notably, the intensity of “free” *CLN3* RNA puncta on the surrounding membrane also increased over time after adding Whi3 protein (Fig. 3A), suggesting that diffuse membrane-bound Whi3 was also capable of forming higher-order assemblies with RNA. However, these RNA clusters displayed slow growth in intensity over time, with a power law scaling exponent of approximately 0.08 ± 0.01, 95% CI of fit (Fig. 3G), substantially slower than the 1/3 scaling exponent expected for liquid-like droplets (*2*). Based on these findings, we hypothesize that RNA clusters on the surrounding membrane may act as “sinks” that arrest condensate assembly by trapping Whi3 proteins within slow-diffusing complexes. In support of this hypothesis, the Whi3 recovery time constant from fluorescence recovery after photobleaching (FRAP) was more than twice as slow in the presence of RNA (Fig. S6E; 221 ± 15 and 97 ± 4 s, 95% CI of fits, with and without RNA, respectively). Once assembled, these Whi3-RNA clusters appear to resist coarsening into macroscopic condensates (Fig. 3G), contrasting with the droplets that form when Whi3 and RNA are mixed in solution (*12, 38*).

These results suggest that membranes are capable of staggering the order of molecular interactions during assembly in a manner that impacts coarsening dynamics. Specifically, homotypic interactions amongst membrane-bound Whi3 proteins presumably favor formation and brief coarsening of macroscopic condensates. However, subsequent, heterotypic interactions between Whi3 and RNA appear to drive a relatively slow clustering process that further restricts coarsening (Fig. 3G). Importantly, this staggering of interactions relies on tethering Whi3 to membranes, followed by recruitment of *CLN3* from solution. We next tested the impact of reordering the interactions on condensate assembly and coarsening dynamics by tethering RNA to membranes rather than protein.

### Membrane-tethered RNA recruits Whi3 and forms clusters that resist coarsening

Based on our finding that recruitment of RNA from solution can limit the coarsening of homotypic, protein-only condensates (Fig. 3 A-C), we hypothesized that even smaller condensates may assemble if Whi3-RNA heterotypic interactions were biased to occur earlier. To test this hypothesis, we directly tethered *CLN3* to membranes by hybridizing RNA to a biotinylated DNA oligonucleotide. Neutravidin protein facilitated RNA tethering to SLBs containing a biotinylated lipid (Fig. 3H). SLBs did not contain Ni-NTA lipid, ensuring that Whi3 recruitment occurred only through RNA binding. Experiments were again performed at physiological ionic strength (150 mM KCl), and Whi3 was added at concentrations far below the saturation concentration for solution LLPS (Fig. 2H) (*12*).

Prior to addition of Whi3, RNA molecules appeared as sparse puncta diffusing rapidly on the membrane at an initial density of approximately 5 puncta/10 µm^2^ (Fig. S7 A-B). Upon addition of 50 nM Whi3, tethered RNA recruited protein from solution and RNA puncta rapidly increased in fluorescence intensity within 30 s (Fig. 3I and S7C; Movie S5). By contrast, when RNA was tethered to an immobile, PEG-biotin surface at a density of approximately 9 puncta/10 µm^2^ (Fig. S7 A-B), RNA only minimally recruited Whi3 and RNA puncta did not increase in fluorescence intensity over time (Fig. 3J and S7C; Movie S6). Thus, Whi3 was capable of rapidly clustering RNA into higher-order assemblies only when RNA was tethered to a diffusive membrane surface (Fig. 3K). Notably, these diffraction-limited *CLN3* clusters (Fig. 3I) were similar in appearance to the *CLN3* clusters assembled by membrane-bound Whi3 (Fig. 3D). Moreover, membrane-tethered *CLN3* clusters grew very slowly over time after the initial clustering, with a power law scaling exponent of 0.05 ± 0.01, 95% CI of fit (Fig. 3L), comparable to the slow clustering of RNA recruited by membrane-bound Whi3 (Fig. 3G). Thus, heterotypic Whi3-RNA assemblies appear to consistently resist coarsening into macroscopic condensates at membranes, regardless of whether Whi3 or *CLN3* was membrane-tethered.

Importantly, these punctate condensates resemble the appearance of Whi3 assemblies in living *Ashbya* cells (Fig. 1A). We found that the scaled distributions of Whi3 puncta intensities from live cells and from reconstitution experiments were both right-skewed in shape and showed substantial overlap (Fig. 3M). This kind of size distribution is expected for Brownian motion coalescence (BMC), in which condensates grow by colliding and fusing (*2*). However, the power law scaling exponent of membrane-tethered *CLN3* clusters was substantially slower than the expected BMC dynamic scaling exponent of 1/3 (Fig. 3L) (*2, 8*). Therefore, while BMC may have possibly contributed to the assembly of Whi3-RNA clusters, another mechanism likely suppressed coarsening. What is the physical mechanism by which Whi3-RNA clusters resist coarsening at membrane surfaces?

### Membrane-tethered RNAs cluster by diffusion-limited aggregation

We wondered if membrane-associated clusters may increase in size and eventually form macroscopic condensates if the concentration of Whi3 in solution increased. We therefore performed membrane-tethered *CLN3* experiments over a range of bulk Whi3 concentrations, from 20-800 nM (Fig. 4 A-B). Surprisingly, Whi3 consistently formed punctate Whi3-RNA clusters at all concentrations, which remained diffraction-limited and did not coarsen into macroscopic condensates (Fig. 4A). Specifically, Whi3 drove rapid RNA clustering within 30 s of protein addition, but clusters quickly plateaued at a relatively constant final puncta intensity (Fig. 4B). However, the intensity of both *CLN3* and Whi3 within clusters increased monotonically as a function of Whi3 concentration (Fig. 4C and S7D), indicating that clusters incorporated more molecules as more Whi3 was available in solution. As expected, we did not observe substantial clustering when RNA was tethered to immobile, PEG-biotin surfaces and the Whi3 concentration ranged from 50-400 nM (Fig. 4C). Analysis of the ratio of Whi3 per *CLN3* in membrane-associated clusters revealed a Langmuir-like trend, approaching an adsorption saturation density of approximately 10 ± 1 Whi3 per *CLN3* (95% CI of fit) (Fig. 4D). This finding suggests that as Whi3 concentration increased, more proteins adsorbed to each RNA (up to approximately 10 proteins), thereby increasing the effective “stickiness” of the RNA and its probability of clustering. Consistent with this hypothesis, RNA also appeared to cluster more rapidly as Whi3 concentration increased (Fig. 4B).

**Fig. 4.**
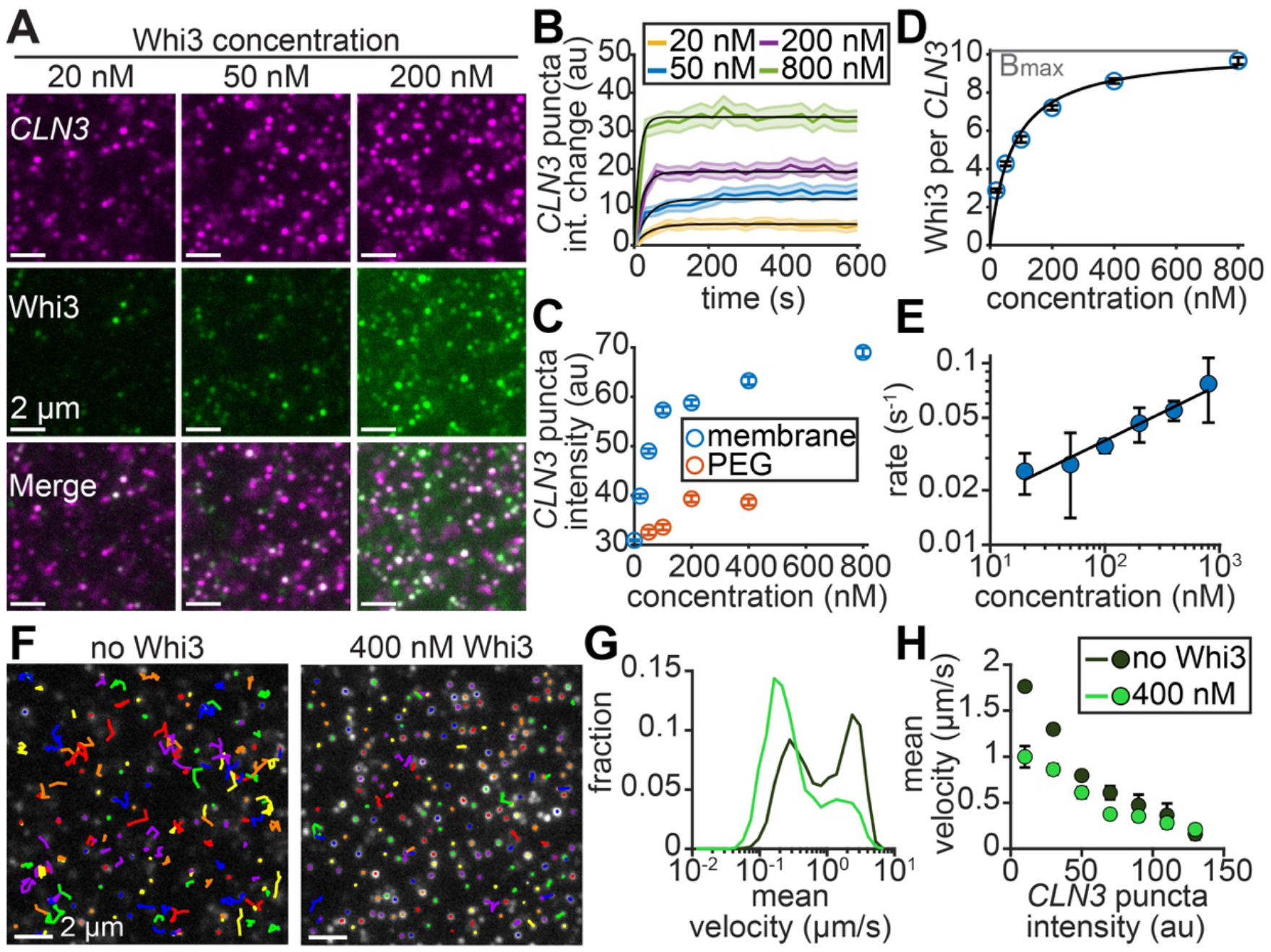
Membrane-tethered *CLN3*-Whi3 clusters assemble by diffusion-limited aggregation. (**A**) Membrane-tethered *CLN3* after 10 min exposure to Whi3 at the indicated concentrations. *CLN3* and Whi3 channels contrasted equally in all images. (**B**) Change in *CLN3* intensity as a function of time after addition of Whi3 at the indicated concentrations. Black lines indicate fits to a single exponential function. (**C**) Average intensity of *CLN3* puncta on membrane or PEG surfaces as a function of Whi3 concentration. (**D**) Ratio of Whi3 per *CLN3* molecule on membranes as a function of Whi3 concentration. Black line indicates fit to the Langmuir isotherm, with fitted Kd and Bmax (gray line) of 72 ± 32 nM and 10 ± 1 Whi3/*CLN3* (95% CIs of fit), respectively. (**E**) *CLN3* clustering rate constant, obtained from fits in (B), as a function of Whi3 concentration. Black line indicates power law fit, with scaling exponent 0.31 ± 0.08 (95% CI of fit). (**F**) RNA puncta tracks before (left) and after (right) addition of 400 nM Whi3. Images contrasted equally. Each color represents a separate track. Tracks corresponding to pinned puncta appear as dots. Puncta were tracked at 100 ms/frame for 3 s, and instantaneous velocities between the positions within each track were calculated. These values were averaged for each track to compute mean velocities shown in (G-H). (**G**) Distributions of mean *CLN3* puncta velocity on SLBs before and after addition of 400 nM Whi3. (**H**) Mean *CLN3* puncta velocity as a function of puncta intensity (measured at the start of the track), before and after addition of 400 nM Whi3. Plot shows moving averages of the raw data, with *CLN3* intensity increments of 20. n > 1,200 puncta per time point for each curve in (B); n > 3,800 puncta per data point in (C); n > 15,000 puncta per data point in (D); n = 9,128 and 3,389 tracked puncta in no Whi3 and 400 nM Whi3 distributions, respectively, in (G); n ranges from 4 - 4,321 puncta per data point in (H). All data pooled from at least three independent experiments. Error bars in (C, D, and H): 95% CI; (E): 95% CIs from fits. SLB membrane composition: 99 mol% DOPC, 1 mol% DOPE cap-biotin.

These observations suggest that RNA clustering by Whi3 may be analogous to the diffusion-limited aggregation of colloids in two dimensions (*45*). Importantly, this mechanism predicts that the colloid clustering rate scales as a power law with the colloid concentration (*46*). Consistent with this prediction, we found that the RNA clustering rate, obtained from fits to a single-component exponential function (Fig. 4B), increased with bulk Whi3 concentration following a power law trend (Fig. 4E). The average intensity of *CLN3* within puncta also scaled with roughly similar power law-like behavior as a function of Whi3 concentration (Fig. 4E and S7E), another prediction of diffusion-limited aggregation (*47*). Together, these results suggest that Whi3 adsorption to *CLN3* transforms RNA into sticky particles that cluster upon collision in a diffusion-limited manner. Notably, this mechanism is conceptually akin to BMC and also yields right-skewed cluster size distributions like those observed for Whi3 puncta in cells (Fig. 3M) (*47, 48*).

Diffusion-limited aggregation implies that clusters should continue to grow over time if diffusion remains unaffected. However, the rapid plateau in cluster intensity within 3-4 min (Fig. 4B) suggests that diffusion may have been hindered. We therefore analyzed the diffusion of membrane-tethered RNA puncta on SLBs before and after addition of Whi3. Notably, the proportion of puncta with average velocity below 0.5 µm/s increased from 39% to 71% upon addition of Whi3 (Fig. 4 F-G), indicating that clustering slowed or effectively stopped RNA diffusion. We also found that prior to clustering, RNA puncta diffusion was inversely correlated with puncta intensity (Fig. 4H), suggesting that puncta containing more RNA molecules likely experienced greater drag and diffused more slowly. However, the average diffusion of a given RNA puncta size was consistently slower in the presence of Whi3 compared to protein-free puncta (Fig. 4H), suggesting that Whi3 simply binding to RNA was also capable of slowing RNA diffusion. Together, these findings reveal that Whi3 binding and subsequent clustering dramatically slows *CLN3* diffusion. This negative feedback between clustering and diffusion likely reduces the probability of collisions with new molecules, rapidly halting cluster growth. These results could explain why clusters remain diffraction-limited in size and do not coarsen into macroscopic condensates.

## Discussion

The mechanisms by which cells localize condensates and control their sizes have remained relatively unexplored. Whi3-RNA condensates function by positioning cell cycle and cell polarity signaling to distinct regions in the cell, indicating that condensate localization is intimately tied to function (*12, 36-39*). Our finding that Whi3 assemblies persistently associate with the ER suggests that attachment to endomembranes plays a key role in controlling condensate localization. The stable, punctate sizes of these condensates prompted us to examine how lipid bilayers may inform Whi3-RNA condensate assembly. Here we discovered that membrane surfaces can generate a stable population of relatively small condensates, which mimic features of *in vivo* condensates, by reducing the diffusive mobility of proteins and RNA (Fig. 5). Specifically, our findings suggest a tradeoff between a local enhancement in protein concentration at membrane surfaces, which favors condensation, and a simultaneous reduction in diffusional mobility, which restricts coarsening. The relative reduction in diffusion appears to determine the condensate size distribution (Fig. 5D).

**Fig. 5.**
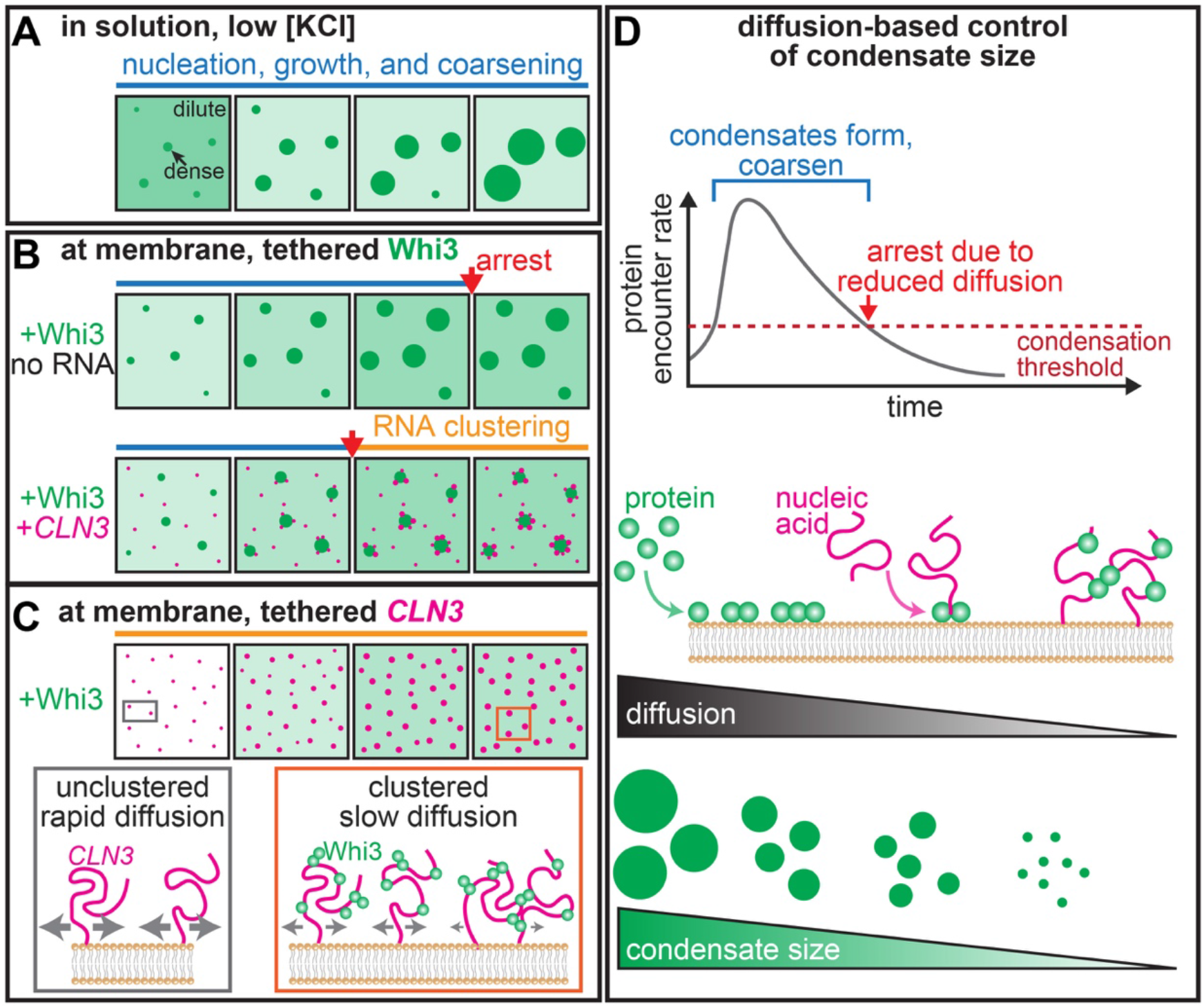
Mechanisms of Whi3-RNA condensate assembly and coarsening at membrane surfaces. (**A**) Whi3 protein-only condensates assembled in solution nucleate, grow, and coarsen into micrometer-scale droplets over several hours (blue regime). (**B**) Top: Recruitment of Whi3 to membrane surfaces drives rapid assembly of micrometer-scale condensates that arrest (red arrow) within minutes of appearance at smaller sizes compared to solution droplets. Bottom: In the presence of RNA recruited from solution, coarsening is arrested even faster, but a subsequent RNA clustering regime at condensate edges continues (orange regime). (**C**) Addition of Whi3 to membrane-tethered *CLN3* results in punctate clusters that assemble via diffusion-limited aggregation and do not coarsen. Boxes: Lateral diffusion of membrane-tethered RNA is hindered upon Whi3 binding and clustering. Sizes of gray arrows indicate relative magnitudes of diffusion. Diffusion-limited aggregation likely governs RNA clustering regardless of whether Whi3 or *CLN3* is membrane-tethered. (**D**) Top: Proposed mechanism for condensate assembly and arrest with membrane-tethered proteins. Recruitment to the membrane concentrates protein, increasing the encounter rate above a condensation threshold. The subsequent reduction in diffusion due to oligomerization in the dilute phase reduces the encounter rate below the threshold, arresting coarsening. Bottom: The magnitude of the reduction in protein and nucleic acid diffusion at membrane surfaces controls condensate size.

In the absence of RNA, recruitment to a membrane locally concentrates Whi3 by more than three orders of magnitude and drives the formation of macroscopic condensates via homotypic protein interactions, consistent with previous findings (*9, 18, 24*). However, the slower diffusion compared to free-diffusing protein in solution results in a faster arrest of coarsening compared to solution droplets (Fig. 5 A, B, and D). When RNA is recruited from solution by membrane-bound Whi3, protein diffusion becomes even more restricted due to the greater hydrodynamic drag of large, membrane-bound Whi3/RNA complexes compared to Whi3 alone. This rollover from primarily homotypic to heterotypic interactions results in a faster arrest and a smaller size distribution compared to protein-only condensates (Fig. 5 B and D). When heterotypic interactions are biased to occur earlier, by recruiting Whi3 to membrane-tethered RNA, Whi3 diffusion is immediately restricted to such an extent that macroscopic condensates do not form at all, but instead remain as small puncta that resemble native assemblies in cells (Fig. 1A and 5 C-D). These results suggest that membrane-bound Whi3/RNA assemblies may inherently resist coarsening into macroscopic condensates simply due to the diffusion constraints experienced by large protein-RNA complexes on membrane surfaces. Notably, this mechanism of size control contrasts with the view that non-equilibrium processes are required to maintain an emulsion of small condensates (*49-51*). Rather, membrane surfaces can offer control over condensate size, without contributions from active mechanisms, by controlling diffusion and the temporal ordering of homotypic and heterotypic interactions.

Is a reduction in diffusion sufficient to explain the hindered coarsening dynamics we observe? To address this question, we computed the approximate rates of diffusion-limited encounter among proteins in solution or at membranes using FCS-based measurements of Whi3 diffusivity (Fig. 2L). Given the likely transient nature of Whi3 homotypic interactions, we hypothesized that Whi3 may require a relatively high encounter rate in order to phase separate. At 50 nM in solution, we estimate an encounter rate of 5 ms, which is presumably prohibitive for phase separation as Whi3 alone does not form solution droplets at this concentration. In SLB experiments, the high local density of Whi3 at the time of condensate appearance (approximately 80 µM, Fig. S4C) suggests that the encounter rate was driven above a condensation threshold (Fig. 5D). However, subsequent Whi3 oligomerization in the dilute phase on the membrane may have progressively slowed protein diffusion and encounter rate back down below this threshold, leading to an arrest of coarsening (Fig. 5D). Consistent with this hypothesis, we estimate that the encounter rate at the approximate time of condensate arrest was 10 ms (Fig. S4C; see methods), comparable to the “prohibitive” encounter rate for dilute proteins in solution. Notably, this reduction in diffusion-limited encounter is likely even more pronounced for membrane-tethered *CLN3* RNAs, which frequently became pinned or immobile after clustering by Whi3 (Fig. 4F). The stalled diffusion of these *CLN3*/Whi3 clusters likely reduced the encounter rate to such an extent that these assemblies never coarsened into macroscopic droplets.

These estimates suggest that Whi3 oligomerization in the dilute phase may play a critical role in controlling the driving forces for phase separation on membranes. How can small oligomers arrest coarsening? Recent work found that condensates can become kinetically arrested in a metastable state if the molecules within small clusters or oligomers can rearrange and self-interact on a faster timescale compared to the encounter rate with new molecules (*52*). We speculate that the Q-rich region of Whi3 may have a finite (although unknown) valency, and that Whi3 condensates may be prone to kinetic arrest if protein diffusion is sufficiently reduced. Specifically, slow-diffusing oligomers on membranes may have time to self-interact and become saturated before encountering condensates, thus becoming “inert” particles that kinetically arrest coarsening. This population of inert particles with satisfied valencies may increase as Whi3 forms clusters with RNA that further slow protein diffusion, thereby driving kinetic arrest even faster. Thus, membranes appear to be capable of controlling condensate size by tuning the extent of protein oligomerization in the dilute phase. These oligomers may act as off-pathway assemblies that negatively regulate coarsening. In support of this hypothesis, our group’s recent work found that a transient ordered domain within Whi3 is also capable of driving dilute phase oligomerization that negatively regulates phase separation in solution (*53*).

Membrane surfaces may also suppress coarsening by driving structural changes within the dense phase. Specifically, the dramatic shift in the phase boundary when Whi3 is recruited to membranes may be analogous to the concept of polyphasic linkage, in which ligand interactions with phase-separating proteins modulate the driving forces for condensation (*54*). From this perspective, membrane surfaces act as “ligands” that bind Whi3 and stabilize condensates under conditions that are otherwise unfavorable for phase separation. However, this additional binding interaction may also slow the exchange of Whi3 proteins between the dense and dilute phases, accelerating the formation of a system-spanning, viscoelastic protein network within the dense phase that further arrests coarsening. Consistent with this hypothesis, the relatively low mobile fraction of Whi3 proteins measured by FRAP (Fig. S6E) was similar to previous measurements of aged, gel-like Whi3 droplets (*12*).

Finally, RNA molecules like *CLN3* that are substantially larger compared to their protein binding partners (*38*) may require considerable conformational freedom in order to build a multivalent network and form macroscopic droplets. In solution, RNA molecules can likely explore a variety of conformations that enable droplet coarsening (*12, 38*). However, anchoring to a membrane surface likely incurs an entropic penalty and restricts RNA conformational freedom, potentially contributing to the kinetic arrest of Whi3/*CLN3* assemblies as small clusters. Importantly, the diffraction-limited clusters observed in tethered RNA experiments (Fig. 3I) resemble the punctate Whi3 condensates seen in living cells (Fig. 1A), suggesting that Whi3 recruitment to ER membrane-tethered RNAs may be a relevant mechanism for condensate assembly *in vivo*. In support of this hypothesis, previous work found that a fraction of virtually all mRNA transcripts associate with the ER for translation, regardless of the type of protein encoded, and that the majority of protein synthesis on the ER is directed toward cytosolic proteins (*55*). Therefore, we hypothesize that a subset of *CLN3* RNA is likely tethered to the ER and may play a key role in recruiting Whi3 and controlling condensate assembly. ER-tethered RNAs may also help control the size and localization of other condensates, including P-bodies and stress granules (*16*).

Biological surfaces such as membranes (*22, 25, 30, 32*), microtubules (*56, 57*), long noncoding RNA (*15*), and DNA (*58-61*) are central features of biological phase separation, but the precise roles of these large surfaces remain unclear. We speculate that the reduction and tuning of protein diffusion via attachment to a biological surface may serve a general function in condensate size control throughout the cell.

## Materials and Methods

### Chemical reagents

HEPES, TCEP, β-mercaptoethanol, and Atto488 NHS-ester were purchased from Sigma-Aldrich. KCl was purchased from VWR. IPTG, lysozyme, imidazole, EDTA, and Pierce protease inhibitor tablets were purchased from Thermo Fisher Scientific. The lipids 1,2-dioleoyl-sn-glycero-3-phosphocholine (DOPC), 1,2-dioleoyl-sn-glycero-3-[(N-(5-amino-1-carboxypentyl)iminodiacetic acid)succinyl], nickel salt (DGS NTA-Ni), and 1,2-dioleoyl-sn-glycero-3-phosphoethanolamine-N-(cap biotinyl) (DOPE cap-biotin) were purchased from Avanti Polar Lipids.

### Plasmids

Full-length *A. gossypii* Whi3 was cloned into a modified version of the pET-30b bacterial expression vector, resulting in a construct with an N-terminal 6his tag followed by an ‘SSG’ linker and a TEV protease cleavage site (although the 6his tag was not removed by TEV cleavage for experiments in this manuscript). The modified pET-30b vector was created by digesting with NdeI and SalI to remove the original purification tags and cleavage sites. A DNA oligo duplex was made by annealing and phosphorylating oligos encoding the 6his tag, ‘SSG’ linker, and TEV site. This duplex was ligated into the digested pET-30b vector to create the modified plasmid backbone. The Whi3 coding sequence was finally inserted into this vector using NotI and SalI restriction sites.

The *A. gossypii CLN3* sequence used for *in vitro* transcription was cloned into the pJet vector (ThermoFisher Scientific K1231) using blunt end cloning. Directionality and sequence were confirmed by Sanger sequencing (Genewiz).

### Protein purification and labeling

Whi3 was expressed as an N-terminal 6his-fusion construct in BL21 *E. coli* cells following induction with 0.5-1 mM IPTG for 16 h at 18 °C. Cells were harvested by centrifugation, and bacteria were lysed using lysis buffer and probe sonication. Lysis buffer consisted of 50 mM HEPES pH 7.4, 1.5 M KCl, 20 mM imidazole, 5 mM β-mercaptoethanol, 1x Pierce protease inhibitor cocktail, and 0.3 mg/mL lysozyme. Protein was purified from crude bacterial extract by incubating with cobalt resin, followed by extensive washing (at least 10x column volumes). Protein was eluted with 220 mM imidazole in 50 mM HEPES pH 7.4, 150 mM KCl, and 5 mM β-mercaptoethanol. Protein was dialyzed overnight against the same buffer without imidazole. Protein concentration was determined by 280 nm absorbance, and stored as small aliquots at -80 °C.

Whi3 was labeled with amine-reactive, NHS ester-functionalized Atto488 (Atto-Tec) in 50 mM HEPES pH 7.4, 150 mM KCl, 5 mM β-mercaptoethanol. The dye concentration was adjusted experimentally to achieve the desired labeling ratio of 0.5-1 dye molecules per protein, typically 2-5 times molar excess of dye. Reactions were performed for 20-30 min at room temperature, and unconjugated dye was removed from labeled protein by overnight dialysis against the above buffer. Before all experiments, protein was centrifuged at approximately 16,000 rcf for 5 min to remove large aggregates.

### *In vitro* transcription

*In vitro* transcription of *CLN3* RNA was carried out according to our established protocols (*38*). The pJet *CLN3* plasmid was linearized using PCR and gel purified (QIAGEN 28706) to create the template for *in vitro* transcription. The template sequence is provided in the Supplemental Information. *In vitro* transcription (NEB E2040S) was carried out according to the manufacturer’s instructions using 100 ng of DNA template. 0.1-1 µL of Cy3-labeled UTP (Sigma PA53026) was included in the reaction to create fluorescently-labeled RNA. Following incubation at 37 °C for 18 h, the reaction was treated with DNAseI (NEB M0303L) according to the manufacturer’s instructions and purified with 2.5 M LiCl precipitation. RNA concentration was determined by 260 nm absorbance and verified for purity and size using a denaturing agarose gel and Millenium RNA ladder (ThermoFisher Scientific AM7151).

In order to tether RNA to biotinylated membranes via neutravidin, the 3’ end of *CLN3* was hybridized to a biotinylated DNA oligo (Integrated DNA Technologies) with the following sequence: 5’ biotin-AAAAAAAAAACAGCTCCAGCGCCTGCACCGCGTAGTTTCT 3’. 10 nM of *CLN3* RNA was mixed with 10 nM of oligo, melted at 95 °C for 5 min, then cooled to 72 °C for 10 min to hybridize.

### Fluorescence microscopy

Imaging was performed on a Nikon Ti-E stand equipped with a Yokogawa CSU-W1 spinning disc confocal unit, motorized TIRF, a Plan-Apochromat 100x/1.49 NA oil immersion objective, a Prime 95b sCMOS camera (Photometrics) for TIRF, and a Zyla sCMOS camera (Andor) for spinning disc. The spinning disc and TIRF were illuminated with separate 488 and 561 nm laser sources. A Bruker Galvo Miniscanner unit was used to perform FRAP experiments.

### Particle detection using cmeAnalysis software

The openly-available cmeAnalysis software (*40*) was used for particle detection and analysis of puncta intensities throughout this manuscript. This software was used for detection of (i) Whi3 puncta in *Ashbya* cells; (ii) single RNA and protein molecules for calibrated fluorescence intensity measurements; (iii) Whi3 puncta on passivated PEG surfaces; (iv) *CLN3* RNA puncta in membrane-tethered Whi3 experiments; and (v) *CLN3* and Whi3 intensities within clusters in membrane-tethered *CLN3* experiments.

The software fit puncta in a specified “master” channel to a two-dimensional Gaussian function with standard deviation equal to the estimated standard deviation of our microscope’s point-spread function (PSF). The software reported the amplitude, “A,” of the Gaussian fit over the local background intensity, “c.” These A values were accepted as valid if they came from diffraction-limited puncta and were significantly above the local c values. The software also returned the centroid positions of detected puncta and, when applicable, searched for puncta in a corresponding secondary channel using the centroids of the detected puncta in the master channel. The search region in the secondary channel was set to three times the standard deviation of the Gaussian fit to the microscope PSF. If a puncta was detected in the secondary channel, the software fit it to a Gaussian function and reported the “A” value of the Gaussian fit over the local background intensity “c” value. If no puncta was detected, the software computed the c value as the average, local intensity in the secondary channel within the search region.

### Cell culture and imaging

The Whi3-tdTomato *Ashbya* strain, tagged at the endogenous locus under the native promoter, was created in our previous work (*38*). The pRS416 A.g.Sec63-GFP:GEN plasmid was transformed into this parent strain by electroporation to create the strain used in this manuscript. Cells were grown in 10 mL *Ashbya* full media (AFM) under selection of G418 (200 µg/ml) and Clonat (100 µg/ml) in a 125 mL baffled flask shaking at 30°C for 16 h. Cells were collected by centrifugation at 300 rpm for 2 min, AFM was removed, and cells were resuspended in low fluorescence media. Cells were placed on a low fluorescence media gel pad containing 2% agarose embedded in a depression slide and sealed with valap. Time-lapse movies of cells were acquired with spinning disc confocal microscopy at a single image plane. Movies were acquired for 15-20 min, 30 s/frame.

### Analysis of Whi3-ER co-localization from single image frames

The first frames of time-lapse movies were used for analysis of Whi3-ER co-localization. All analysis was performed in Matlab 2019b software. The centroid positions of diffraction-limited puncta of Whi3 assemblies were estimated using cmeAnalysis software. The average, local intensity in the ER channel at each detected Whi3 centroid was then computed using a masking procedure. Specifically, a binary image was created at each detected position and dilated into a disc with radius equal to two times the standard deviation of the microscope PSF. The average intensity of the ER channel within the mask was then computed after subtraction of the camera background. We defined the threshold for co-localization as the median intensity of the ER channel throughout the cell, after automatically thresholding using the triangle method to create a mask of the cell. We found that this median intensity value corresponded approximately to the average background intensity within the ER channel outside of labeled ER structures, indicating that it was an appropriate choice for the co-localization threshold. Values of the associated intensity within the ER channel at detected puncta were expressed as a ratio of this threshold. Values greater than one in Fig. 1B were defined as co-localized.

A random number generator was used to create randomized positions of the same number of detected Whi3 puncta. Positions were defined as valid if they fell within the cell mask, did not contact the edge of the cell mask when dilated, and were separated by at least two times the standard deviation of the microscope PSF. The average intensity in the ER channel at each random position was computed using the same masking procedure described above. To validate this procedure, 50 different sets of randomized positions were generated for each cell. We used the following procedure to compute the data shown in Fig. 1C:

i. We computed the average of all of the local ER intensities at each detected puncta within a cell.
ii. For a given set of randomized positions, we computed the average of all of the local ER intensities at each random position within the cell.
iii. For each set of randomized positions, we took the ratio of the averages from steps (i) and (ii), resulting in 50 different values. A value greater than one indicates that the local intensity in the ER channel was, on average, greater at detected puncta relative to randomized puncta.
iv. Finally, each data point in Fig. 1C was computed as the average of these 50 different ratio values.

This procedure was repeated with n = 60 cells. The average ratio of 1.62 ± 0.36 s.d. indicates that local intensity in the ER channel associated with detected Whi3 puncta was consistently greater compared to randomized puncta.

### Tracking Whi3 puncta

The centroids of Whi3 puncta at each movie frame were detected using cmeAnalysis software, which then determined the local “c” values within the ER channel at each Whi3 centroid. These local intensity values in the ER channel were then expressed as a fraction of the median intensity of the ER channel throughout the cell, after masking and background subtraction. Values greater than one were defined as co-localized with the ER. Puncta centroids were linked into tracks using freely-available Matlab code from http://site.physics.georgetown.edu/matlab/. The maximum displacement between positions within a track was approximately 0.5 µm, with a maximum allowable gap of two image frames. Only tracks that included four or more image frames were accepted for further analysis. We estimated the fraction of each track’s lifetime spent co-localized with the ER by computing the proportion of frames within each track with relative, local ER intensity greater than one. Fig. 1E shows the relative, local intensity in the ER channel along three example tracks, each of which spends 100% of the lifetime co-localized with the ER. Fig 1F shows a histogram of 769 tracks, binned according to the fraction of track lifetime co-localized with the ER.

### Supported lipid bilayer (SLB) preparation

Lipid films were prepared by mixing lipids in chloroform at the specified molar ratios, drying under a stream of argon, and placing under vacuum for at least 2 h. All membrane compositions are provided in the figure captions. Lipid films were hydrated and resuspended in 50 mM HEPES pH 7.4, 150 mM KCl buffer at a final lipid concentration of 0.5-1 mM. Small unilamellar vesicles (SUVs) were then created by extruding lipid suspensions through 100 nm-pore filters (Whatman) or using 3 × 4 min cycles of probe sonication at 70% power (Branson Sonifier SFX150). Glass coverslips (VWR 48393-230) were treated with oxygen plasma for 15 min to clean the glass and render the surface hydrophilic. Imaging wells were created using silicone gaskets with a 13 mm-diameter hole (Grace Bio-Labs 664170), or gaskets cut from silicone sheets (Grace Bio-Labs 664172) and punched with a 5 mm-diameter hole. SUVs were added to wells at a final lipid concentration of 0.25 mM and allowed to rupture and form an SLB on the glass surface for 20 min at room temperature. The SLB was washed thoroughly, at least six times, with 50 mM HEPES pH 7.4, 150 mM KCl, 2 mM TCEP buffer to remove excess liposomes. After another 15 min period, the SLB was washed thoroughly again to ensure that all excess liposomes were removed from the well and the SLB surface. For SLBs functionalized with DOPE cap-biotin, neutravidin protein was added at a final concentration of 10 µg mL^-1^, incubated for 5 min, and washed repeatedly to remove unbound neutravidin before adding biotinylated *CLN3* RNA.

### Membrane-tethered Whi3 experiments and analysis of membrane-associated Whi3 condensates

Whi3-Atto488 was added to NTA-Ni-functionalized SLBs at the specified final concentrations and allowed to bind and form condensates for 20 min. During this time, images were collected every 30 s using TIRF microscopy. Finally, approximately 10-20 images of condensates on SLBs were acquired within a period of 20-30 min after addition of Whi3. Experiments were repeated with at least three independent SLBs per condition.

Images were cropped to the center 400x400 pixels (center 1/9) of the original images to ensure even illumination across the field of view to be analyzed. Images were processed using a custom ImageJ script that applied an automatic threshold using the triangle method, applied a watershed, and used the “Analyze Particles” tool to find and outline condensate edges. The resulting ROIs were used to measure condensate features, which were saved and imported into Matlab for further analysis. Condensate radii were estimated from measurements of condensate area by assuming a circular shape. To examine the dynamics of average condensate radius over time, each frame from time-lapse movies was analyzed using a similar ImageJ script to the one described above. The centroids of condensates were linked into tracks using code from http://site.physics.georgetown.edu/matlab/. A maximum displacement of 0.37 µm was used, and only tracks that included 10 or more image frames were accepted for further analysis. Average radius was computed from tracked condensates and plotted as a function of time after condensates first appeared.

To confirm that attachment to a membrane surface was required for condensate assembly, Whi3 was added to passivated, PEG-coated surfaces (described in the section “Passivating glass with PEG and PEG-biotin”) at a concentration of 50 nM. A time-lapse TIRF movie was acquired for 20 min as diffraction-limited Whi3 puncta randomly adhered to the PEG surface. Finally, 10 images of Whi3 puncta were acquired after 20 min and cmeAnalysis software was used to estimate the number of Whi3 proteins per puncta, as described in the section “Determining the intensity of single RNA and protein molecules.”

To examine recruitment of *CLN3* RNA from solution by membrane-bound Whi3, cmeAnalysis software was used to detect the intensities of RNA puncta as well as the associated, local average intensity in the Whi3 channel at each detected RNA puncta. RNA puncta were defined as associated with condensed Whi3 “patches” if the associated, local intensity in the Whi3 channel was greater than a threshold value of 1.25*I_WM_, where I_WM_ is the median intensity of the Whi3 channel throughout the image. We found that this threshold value was in good agreement with the average intensity of condensed Whi3 patches at each time point in the movies. RNA puncta that did not meet this criteria were defined as “free” on the surrounding membrane. We computed the average intensity of RNA puncta in each of these categories at each time point to generate plots of RNA clustering over time. RNA puncta intensity in each of these categories was then analyzed 20-30 min after condensate assembly to generate histograms.

### Fluorescence recovery after photobleaching

Condensates on SLBs were bleached with a 488 nm laser and TIRF images were acquired for 10 min. A region of the SLB outside of condensates was also bleached and used for background subtraction. Fluorescence intensity values within bleached regions were exported using ImageJ. Data were normalized to prebleach levels and fitted to a single exponential recovery function, *F*(*t*) = *A*[1 − exp (−*t*/*τ*)], where F(t) is the relative fluorescence intensity over time, A is the mobile fraction, and *τ* is the recovery time constant. Fitting was performed in Matlab 2019b software.

### Passivating glass with PEG and PEG-biotin

Glass coverslips were passivated by coating with poly-L-lysine (PLL) conjugated to polyethylene glycol (PEG) and biotin-PEG, prepared according to previous protocols (*62, 63*). Briefly, amine-reactive PEG-succinimidyl valerate (SVA) and biotin-PEG-SVA was added to a 40 mg mL^-1^ mixture of PLL in 50 mM sodium tetraborate pH 8.5 at a molar ratio of one PEG per five lysine subunits. PEG-biotin comprised ∼2% of the total PEG amount. The mixture was stirred continuously for 6 h at room temperature and buffer exchanged into 50 mM HEPES pH 7.4, 150 mM KCl using Centri-Spin size exclusion columns (Princeton Separations). Imaging wells were made by placing silicone gaskets on oxygen plasma-treated coverslips and coated for 20-30 min with PLL-PEG diluted tenfold in 50 mM HEPES pH 7.4, 150 mM KCl. After coating, the well was washed repeatedly with the same buffer to remove excess PLL-PEG. Neutravidin was added at a final concentration of 10 µg mL^-1^, incubated for 5 min, and washed repeatedly again.

### Surface-tethered *CLN3* experiments and analysis of Whi3/*CLN3* clusters

*CLN3* RNA, hybridized to a biotinylated DNA oligo, was added to wells a final concentration of either 200 pM or 10 pM for biotin SLBs or biotin-PEG surfaces, respectively. RNA was allowed to bind to neutravidin for 15 min, and wells were washed thoroughly to remove unbound RNA. 3-5 images of RNA on SLB or PEG surfaces prior to addition of Whi3 protein were first acquired using TIRF microscopy. Whi3 protein was then added at the specified concentrations, and time-lapse images were acquired every 30 s for 10 min. Finally, approximately 10-20 images of Whi3/*CLN3* clusters were acquired within 10-20 min after addition of Whi3 protein.

Images were cropped to the center 400x400 pixels (center 1/9) of the original images to ensure even illumination across the analyzed field of view. Images were analyzed using cmeAnalysis software with *CLN3* and Whi3 specified as the master and secondary channels, respectively. We estimated the number of *CLN3* and Whi3 molecules within each cluster by comparing the brightness values in each channel to the brightnesses of single molecules of *CLN3*-Cy3 and Whi3-Atto-488, respectively (described in the next section). Finally, these values were used to compute the ratio of Whi3 proteins per RNA within each cluster.

To analyze the velocities of diffusing RNA puncta on membranes, time-lapse movies of RNA before and after addition of 400 nM Whi3 were acquired for 3 s, 100 ms/frame. The RNA puncta were detected using cmeAnalysis software, and particle centroids were linked into tracks using code from http://site.physics.georgetown.edu/matlab/, with maximum displacement 0.59 µm between frames. Tracks spanning at least 4 frames were accepted for further analysis. Track velocities were extracted using the freely-available msdanalyzer package for Matlab (*64*), which provided the instantaneous velocities between positions within each track. These values were averaged over each track to compute mean track velocities.

### Determining the intensity of single RNA and protein molecules

Images of single molecules were acquired by creating imaging wells on oxygen plasma-treated coverslips and adding *CLN3*-Cy3 or Whi3-Atto488 at a concentration of 50 pM. Images of single molecules adhered to the coverslip surface were acquired using the same TIRF and laser power settings used in SLB or PEG surfaces experiments, but with longer exposure times to acquire sufficient signal. Images were cropped to the center 400x400 pixels to ensure even illumination, and the diffraction-limited puncta of single protein molecules were detected using cmeAnalysis software. “A” values were pooled from 5-10 image frames and binned into a histogram, yielding a distribution with one clear peak corresponding to the average intensity of a single molecule. Finally, we estimated the number of molecules per puncta in experiments by comparing to this single molecule intensity, after linearly adjusting for differences in camera exposure time.

This approach was validated for Whi3-Atto488 using single-step photobleaching measurements. Specifically, 100 images of surface-adhered Whi3 proteins were acquired continuously using the same TIRF, laser, and exposure settings, and we identified puncta that bleached completely to the level of the camera background in one, clear step. We found that the average, pre-bleach peak intensities of these puncta were in agreement with our estimates from particle detection. See Fig. S3 for more detail on this workflow.

### Quantifying protein density on membranes

The density of membrane-bound proteins on the surrounding SLB (outside of condensates) was computed using the median protein intensity at the SLB after subtraction of the camera background. Because the SLB surface was densely covered by labeled Whi3 proteins, with relatively uniform intensity throughout, the number of labeled molecules, N, per surface area, a, is given by

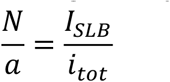

where I_SLB_ is the uniform intensity of the densely labeled SLB and i_tot_ is the total integrated intensity of a single molecule. However, estimates of single protein intensity obtained in the previous section correspond roughly to the integrated intensity within the full width at half maximum (FWHM) of the 2D Gaussian intensity pattern, rather than the total integrated intensity. We call this value i_FWHM_. Therefore, a correction factor must be applied in order to compute the number of molecules per surface area using i_FWHM_. This correction factor can be obtained by considering the 2D Gaussian integral in polar form.

After evaluating the angular component from 0 to 2*π*, the normalized integral is

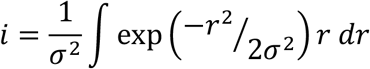

where *σ* is the Gaussian PSF standard deviation. When r is evaluated from 0 to ∞, the total integrated volume is i_tot_ = 1. When r is evaluated from 0 to 1.18*s, corresponding to the Gaussian FWHM, the integrated volume is i_FWHM_ = 0.5, indicating that the FWHM integrated intensity of a single molecule is 1/2 of the total integrated intensity. Therefore, *i*_*tot*_ *=* 2 *i*_*FWHM*_, and the number of labeled molecules per surface area is

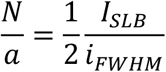

where a is the Gaussian PSF FWHM area. Finally, the total number of protein molecules per surface area was determined by adjusting for the known fraction of dye-labeled to unlabeled protein.

We estimated that the density of membrane-bound Whi3 proteins was approximately 217 proteins/µm^2^ when condensates first appeared. To estimate the pseudo-3D protein concentration corresponding to this density, we assumed that proteins occupied a volume with z-dimension equal to the average diameter of Whi3. This diameter was taken as twice the hydrodynamic radius, *R*_*H*_, of Whi3, estimated from FCS to be 2.3 ± 0.4 nm s.d. (see section “Fluorescence correlation spectroscopy”). This yielded an approximate pseudo-3D concentration of 78 µM.

### Assembly and analysis of droplets in solution

Whi3 was concentrated using a 30 kDa molecular weight cutoff centrifugal filter (Amicon) and diluted into 50 mM HEPES pH 7.4 and 2 mM TCEP buffer at the specified protein concentration such that the final KCl concentration was 75 mM. Atto488-labeled Whi3 protein was included such that approximately 0.1% of Whi3 was dye-labeled. Protein solutions were loaded into glass-bottom imaging chambers (Grace Bio-Labs). To prevent protein adsorption to the imaging surface, wells were passivated with PLL-PEG and washed thoroughly with experiment buffer prior to addition of Whi3 protein solutions. Confocal z-stacks of droplets were acquired at the specified times after droplet assembly, starting below the coverslip surface and with a z-spacing of 0.2 µm. Three z-stacks were acquired at each time point, and experiments were replicated three times. Maximum intensity projections were created from z-stacks, and images were automatically thresholded using the Otsu method and analyzed to obtain values of droplet projected area. Finally, droplet radii were calculated by assuming a circular projected area.

### Fluorescence correlation spectroscopy

To ensure that all unconjugated dye was removed prior to fluorescence correlation spectroscopy (FCS) experiments, Whi3-Atto488 was buffer exchanged into 50 mM HEPES pH 7.4, 2 mM TCEP buffer containing 1 M KCl using a CentriSpin-20 size exclusion column (Princeton Separations), followed by a second buffer exchange back into 50 mM HEPES pH 7.4, 2 mM TCEP, 150 mM KCl buffer. Protein was then ultra-centrifuged for 10 min at approximately 200,000 rcf_avg_ to remove large aggregates.

For measurements of protein diffusion at membranes, NTA-Ni-functionalized SLBs were prepared as described above and washed with 50 mM HEPES pH 7.4, 150 mM KCl, and 2 mM TCEP buffer. Whi3-Atto488 was then added at a concentration of 2 nM and allowed to bind the membrane for 20 min. The SLB was washed thoroughly to remove all Atto488-labeled protein from solution. Finally, unlabeled Whi3 protein was added to the well at a final concentration of 50 nM in order to match the conditions used for assembly of membrane-associated Whi3 condensates. For measurements of protein diffusion in solution, the glass surface was passivated with a pure DOPC SLB to prevent protein adsorption, and Whi3 was added to wells at a concentration of 50 nM. Whi3-Atto488 was included in protein mixtures such that the final Atto488 dye concentration was 3-6 nM. Calibration standards of Atto488 dye alone and GFP at concentrations of 1 and 2 nM, respectively, were prepared in similar wells. For membrane-bound GFP experiments, 2 nM GFP (which contained an N-terminal 6his tag) was added to NTA-Ni-functionalized SLBs, allowed to bind for 20 min, and washed thoroughly to remove unbound protein.

FCS traces were acquired on a Zeiss LSM 880 microscope using a 40x water immersion objective. To measure protein diffusion at the membrane or free in solution, a 488 nm laser was either focused at the coverslip surface or 200 µm into solution, respectively. Fluorescence signal was collected with a Zeiss BiG.2 GaAsP detector. Each FCS trace was acquired for 100 s and fit with the 2D, single-component autocorrelation function,

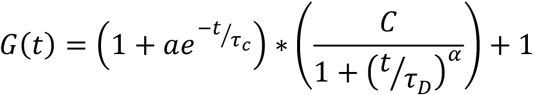

where *C* is 1/*N*_*p*_, *N*_*p*_ is the number of labeled proteins in the focused laser volume, *τ*_*D*_ is the diffusion time, and *α* is the anomalous diffusion coefficient. *a* and *τ*_*c*_, which correct for short time processes such as intersystem crossing, were held constant in the fitting as 0.05 and 5 µs, respectively (*43*). Fitting was performed in Matlab 2019b software.

Using the measured *τ*_*D*_ of Atto488 dye, with well-characterized diffusion coefficient, *D*, of 400 µm^2^/s, we computed the lateral radius of the focal volume, *r*_0_, by the relationship 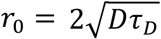. Using this estimated *r*_0_, we computed the diffusion coefficients of Whi3 and GFP in solution or at membranes. To check the validity of our approach, we estimated the hydrodynamic radius, *R*_*H*_, of GFP in solution using the Stokes-Einstein relationship,

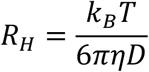

where *k*_*B*_ is the Boltzmann constant, *T* is the absolute temperature, and *η* is the dynamic viscosity of water at room temperature. This yielded an estimated *R*_*H*_ for GFP of 2.2 ± 0.6 nm s.d., in reasonable agreement with a previous estimate of 2.5 nm (*65*).

Importantly, FCS measurements of membrane-bound Whi3 were performed in the “dilute” phase, on the surrounding membrane and away from Whi3 condensates. This was achieved by imaging the SLB prior to acquisition, and checking that the average fluorescence count rate at the start of acquisition was consistent among traces.

Measurements were taken at least 20 min or more after addition of 50 nM Whi3 to wells. Diffusion coefficient estimates therefore reflect the diffusivity of membrane-bound Whi3 after the onset of condensate arrest.

### Estimation of Whi3 encounter rates

The approximate rate of Whi3 encounter in solution was determined by first estimating the average spacing among Whi3 proteins at a concentration of 50 nM. Assuming equidistant spacing, the average volume around each protein is approximately 0.033 µm^3^, corresponding to a surface area surrounding each protein of 0.5 µm^2^ (assuming a spherical volume around each protein). Based on our FCS-based estimate of the Whi3 diffusion coefficient in solution of 108 µm^2^/s, proteins will explore this surface area within approximately 5 ms.

The approximate encounter rate of membrane-bound Whi3 proteins at the time of condensate arrest was estimated using the measured Whi3 protein density on the surrounding SLB 4.5 min after addition of Whi3 (Fig. S4C, red lines). This was the approximate time at which condensate radii reached a plateau, and corresponded to a membrane-bound protein density of 563 proteins/µm^2^. At this density, the average area around each protein is approximately 0.0018 µm^2^ (assuming equidistant protein spacing). Assuming that the average diffusion coefficient of membrane-bound Whi3 proteins was 0.17 µm^2^/s (measured from FCS) at the time of condensate arrest, Whi3 proteins will explore this area within approximately 10 ms.

## Acknowledgements

A.S.G. acknowledges funding from the National Institutes of Health (R01-GM081506), the Howard Hughes Medical Institute (HHMI) Faculty Scholars program, and the Air Force Office of Scientific Research (FA9550-20-1-0241). W.T.S. acknowledges the support of a Ruth L. Kirschstein NRSA Postdoctoral Fellowship from the National Institutes of Health (F32-GM136055). We thank Dr. Christine Roden of the Gladfelter Lab for cloning the *CLN3* plasmid and performing *in vitro* RNA transcription. We thank Dr. Carl Hayden (UT Austin, Austin, TX) for helpful discussions on image analysis. We thank Tony Perdue and the UNC Biology Department Imaging Core for assistance with FCS and use of the Zeiss LSM 880 microscope. We thank Dr. Gaudenz Danuser and Dr. Sandra Schmid (UT Southwestern, Dallas, TX) for freely providing cmeAnalysis particle detection software.

The authors declare no competing financial interests.

## Author contributions

All authors designed and performed experiments, consulted together on the data, and wrote the manuscript.

