## Supplemental Information for "Membrane surfaces regulate assembly of a ribonucleoprotein condensate"

**This file contains**

- Sequence of *CLN3* template used for *in vitro* transcription
- Figs. S1-S7
- Movies S1-S6 captions
- SI References

**Sequence of *CLN3* template used for *in vitro* transcription**

taatacgactcactataggggtctgcataccaaggatcagccgcttgcattaaaggggacgaaccggggctcttatcctta
gcatcgtctcttgtccacacctcgttgacctgcagacacagcacaaaccctgcataattaatagtaatcattgtcatctccag
ctgggctgttaatatctcatacccgaacacctttgcatattatacacatattcgtcatggactctacaaaactctcatgcgataa
ggtttcgctgctacaactggcctcaatcaggaggtctgtgtgggacagtaaggcatcccatccccggttaatacacatgga
gatggttgcgcaccaatctgcctgccaggagtacagcattgaaatcctcaagcatctgattgatctcgagaatcagacccg
gtcgtcatggactagcttccagtcccagccggaacttaccgtggagatgcgcactttgattttcgactttatcatgagctgcca
cactcgcttgggactttcctcctcgacactcttcttatgctacaacatcattgatcggtactgctccaagatcattgttaaatcac
ccacatatcagctactagggcttaccgctctttggctggcgtccaagttcgcagacaaaaagccccgaataccgagcctg
cagagcctctgctgccaccagcacacgaaacagcagttcaaggaaatggagctgcacatcctcaagagccttaactggt
ctgtgtgctctgctccgagtcacgattcgttcgtggatatcctcctgaaaaccaagatagccaatctcagcttcaagcgcctc
aatctaaacgatttaaagtatggagcaacgattctttgcgagctatcctgctttgacccagcactgaactacaattacaactc
cagcgccattgccttggcaagcgttacagttataacctgcgcactccgactcgctgagctctccgagttcatagattgcagg
cgatgtactactgacaaaaatctcattgtcatctgcaaccagcttctctgtctactagcctgcgaggacaacttcccgtccag
ctttaacctcaagtacgcttccgagggcggcacgctccactccaaccccattatggcccgtctgttcgcttattaccgccggtt
agctgatgagaaaaaacgttcgtggcgtgatgcaggcactcgcgtgggctcgccatgtgcaggctcagcaccggccagt
gctgccggcggctcccccgatatccgcattggcggcgacacgcccccgggctctgttccgccttccgctctgggcccgctt
gccccgcatatgccgacaatcaagacttttgtgcctcccacaccaactactccgagcaactcatcgcgcactcagctgtcg
ctttgtgccaagccccatacgctacccgcgcctgtgtccgtcgcagcgtcggcgcgcgcaaaaggcactgcacatcagc
acacgcctccggtccagaaccccgcgcacttgctgcatactgcgcaatctaagcgcgccgcctcggccatggaccttga
gttcttcgacatggatcatgccctggtcaagaaactacgcggtgcaggcgctggagctg

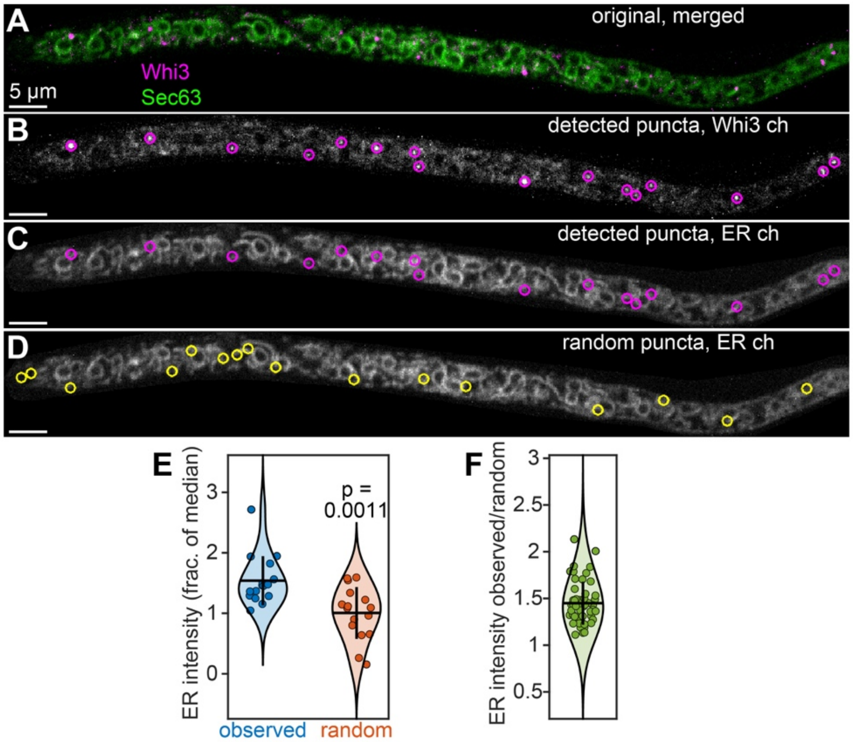


**Figure S1**. Quantifying ER co-localization with detected Whi3 puncta and randomized puncta. (**A-D**) Images of an *Ashbya* hypha expressing Whi3-tdTomato (tagged endogeneously) and ER marker Sec63-GFP (plasmid expression). Images show merged channels (A), detected Whi3 puncta after particle detection (B), the same detected puncta overlaid with the ER channel (C), and a random distribution of the same number of puncta overlaid with the ER channel (D). (**E**) Local intensity in the ER channel at the Whi3 puncta shown in the above images, expressed as a fraction of the median intensity of the ER channel throughout the hypha after masking and background subtraction. Blue points: observed Whi3 positions, orange points: randomized Whi3 positions corresponding to the puncta in image (D). n = 16 puncta. Dashed line indicates the threshold for co-localization, corresponding to the median intensity of the ER channel. 100% of detected puncta are co-localized with the ER, compared to 56% of randomized puncta. Indicated p-value from two-tailed, unpaired Student's t-test. (**F**) Ratio of the average local intensity in the ER channel at detected Whi3 puncta within the above hypha relative to randomized Whi3 puncta positions. A value greater than one indicates that the local intensity within the ER channel is greater on average at detected Whi3 puncta compared to randomized puncta. Each data point represents the ratio to one of 50 random distributions. The mean, indicated by the black horizontal bar, corresponds to one of the 60 data points in Fig. 1C. Vertical bars represent standard deviation.

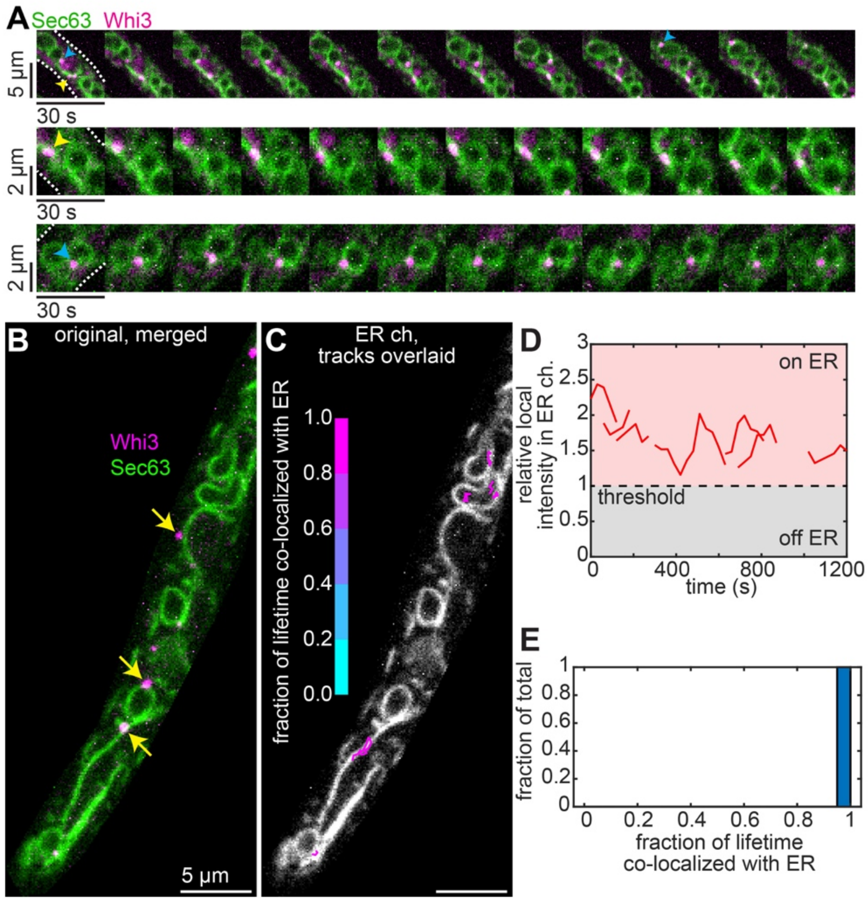


**Figure S2**. Tracking Whi3 puncta and ER co-localization. (**A**) Time-lapse montages of Whi3 puncta associated with ER, including ER tubules (yellow arrowheads) and nuclear-associated ER (blue arrowheads). White dashed lines in first frame indicate cell periphery. Similar to Fig. 1D. (**B**) First frame from time-lapse shown in Movie S1. Yellow arrows indicate puncta that appear co-localized with the ER but moved out of the imaging plane during the movie and were not included in the tracking. (**C**) ER channel from the image in (B) with overlaid Whi3 tracks, colored according to the fraction of the track lifetime spent co-localized with the ER. All tracks clearly co-localize with ER structures. Not all tracks begin in the indicated frame. (**D**) Relative, local intensity in the ER channel as a function of time for the Whi3 tracks shown in (C), expressed as a fraction of the median intensity in the ER channel throughout the cell. Values greater than one (red region) were defined as co-localized with the ER. In this example, all tracks spend 100% of the lifetime co-localized with the ER. (**E**) Histogram of the tracks in (C-D), binned according to the fraction of track lifetimes co-localized with the ER (similar to Fig. 1F). n = 7 tracked Whi3 puncta in this example.

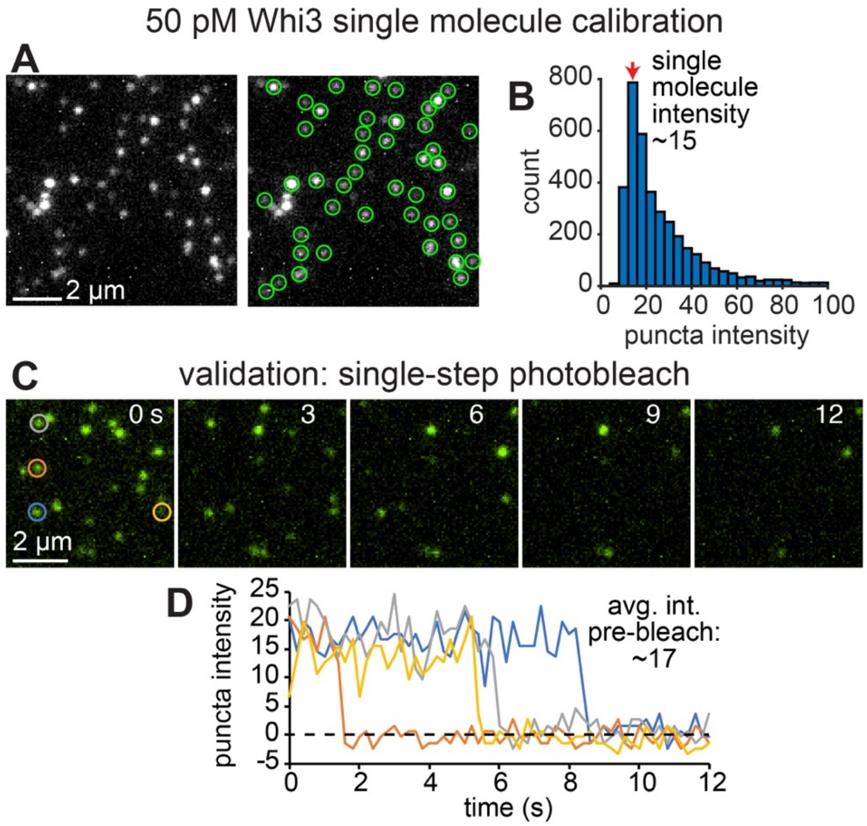


**Figure S3**. Particle detection and single molecule calibration. (**A**) TIRF image of 50 pM Whi3-Atto488 on a plasma-cleaned glass coverslip. Right image shows detected puncta from cmeAnalysis software (*1*). (**B**) Histogram of detected puncta intensities, obtained from fits to a two-dimensional Gaussian function with standard deviation equal to the microscope PSF (see methods). The indicated peak value (red arrow) was taken as the average intensity of a single Whi3 protein. (**C-D**) This single molecule intensity estimate was validated using photobleaching measurements. (C) TIRF time-lapse of 50 pM Whi3-Atto488 puncta on glass at the indicated times. Images were acquired with the same TIRF angle, laser power, and camera exposure settings used for acquisition of single molecule calibration images in (A-B). (D) Average, background-subtracted peak intensities of the puncta in the indicated colored circles in (C). Each puncta bleaches to the level of the camera background in a single step, indicating that each puncta corresponds to a single Whi3-Atto488 protein. The average, pre-bleach intensity of each puncta was comparable to our estimate of the single molecule intensity from particle detection in (B)

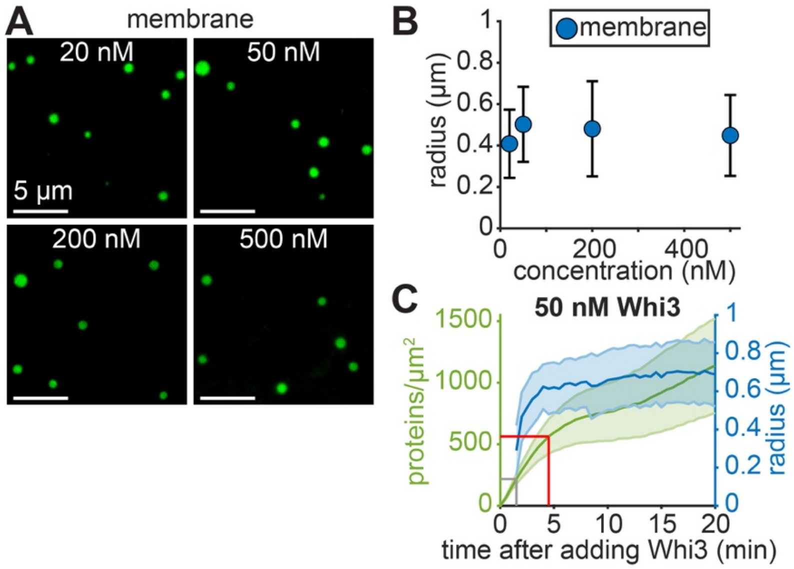


**Figure S4**. Membrane-associated condensate size as a function of bulk protein concentration, and over time compared to the density of membrane-bound protein. (**A**) Images of membrane-associated condensates formed at the indicated bulk Whi3 concentrations, 20-30 min after condensate assembly. (**B**) Membrane-associated condensate radius as a function of bulk Whi3 concentration. n > 250 condensates per data point, pooled from at least three independent experiments. Error bars indicate standard deviation. (**C**) Overlay of average condensate radius (blue, repeated from Fig. 2E) and average protein density on the surrounding SLB (green) as a function of time after adding Whi3. Gray and red vertical lines indicate the approximate times of condensate appearance and arrest, respectively, while gray and red horizontal lines indicate membrane-bound protein densities at those times (217 and 563 proteins/µm$^2$ at condensate appearance and arrest, respectively). Green curve represents the average from n = 3 SLBs. Error bars: standard error of the mean (green curve) and standard deviation (blue curve). SLB membrane composition: 96 mol% DOPC, 4 mol% DGS NTA-Ni.

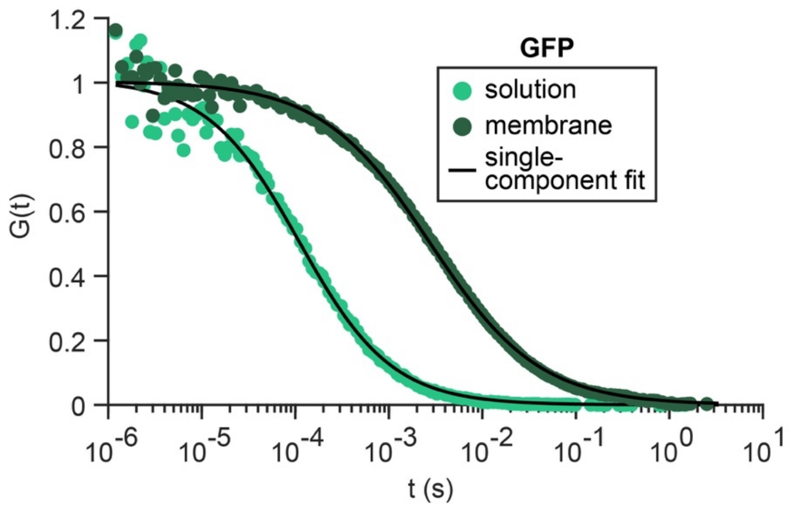


**Figure S5**. Average, normalized FCS traces of GFP in solution or bound to membrane. Black curves indicate fits to single-component diffusion model. See Fig. 2L for estimated GFP diffusion coefficients. SLB membrane composition: 96 mol% DOPC, 4 mol% DGS NTA-Ni.

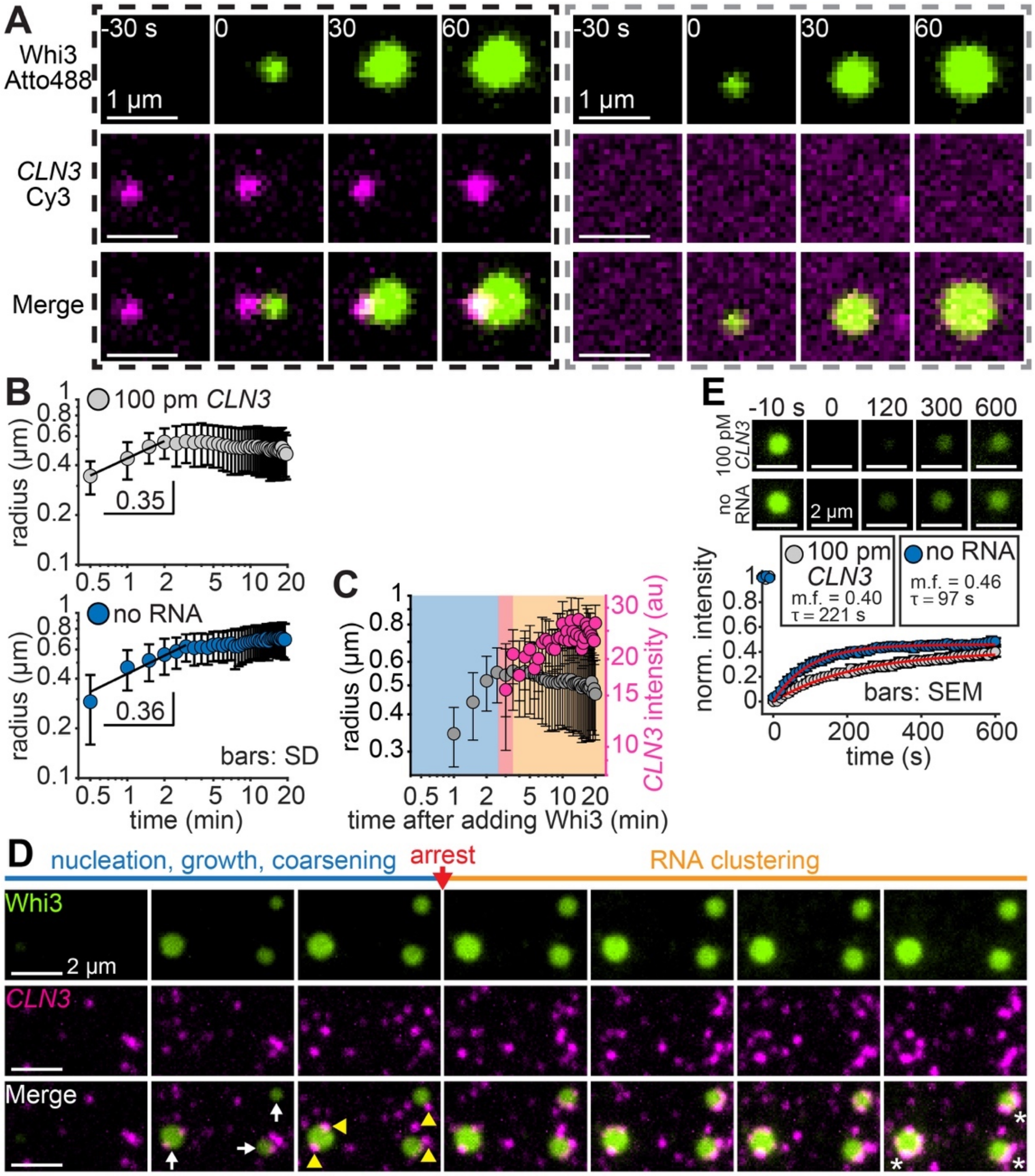


**Figure S6**. RNA suppresses coarsening but does not influence initial Whi3 condensate assembly dynamics. (**A**) Time series of two example condensates formed in the presence of 50 nM Whi3 and 100 pM *CLN3*. Images show condensates formed in proximity to an RNA puncta (black dashed box) and with no associated RNA (gray dashed box). (**B**) Condensate radius in the presence (upper) and absence (lower) of RNA as a function of time after condensates first appear. Black lines show fit to the power law dynamic scaling function $r(t)=(K*t)^n$, where r(t) is the average radius as a function of time, K is the dynamic scaling prefactor, and n is the scaling exponent. Fit is to the first four data points in the presence of RNA (upper) and the first six data points in the absence of RNA (lower), with fitted exponents indicated. The shorter coarsening period in the presence of RNA suggests that RNA may have suppressed coarsening faster compared to RNA-free experiments. n > 50 tracked condensates per condition. Error bars indicate standard deviation. (**C**) Overlay of average condensate radius and average patch-associated *CLN3* puncta intensity. Plot repeated from Fig. 3F, shown here with shaded regions that approximate the coarsening (blue), arrest (red), and RNA clustering (orange) phases. (**D**) Time-lapse of condensate assembly at SLB with 50 nM Whi3 and 100 pM *CLN3*, with approximate coarsening, arrest, and RNA clustering phases indicated by the same colors in (C). Frames span 2.5-8.5 min after addition of Whi3, 1 min between frames. White arrows indicate Whi3 condensates visible after 3.5 min, and yellow arrowheads indicate condensate edge-associated *CLN3* puncta apparent at the approximate time of arrest. White asterisks in final frames indicate bright *CLN3* clusters that continued to assemble after arrest. (**E**) FRAP of condensates formed with 50 nM Whi3 in the presence (upper) and absence (lower) of RNA. Plot shows corresponding FRAP profiles. Red curves show fits to single-component exponential recovery model, with mobile fractions (m.f.) and recovery time constants ($\tau$) indicated. n = 12 and 8 condensates with and without RNA, respectively. Error bars indicate standard error of the mean. SLB membrane composition: 96 mol% DOPC, 4 mol% DGS NTA-Ni.

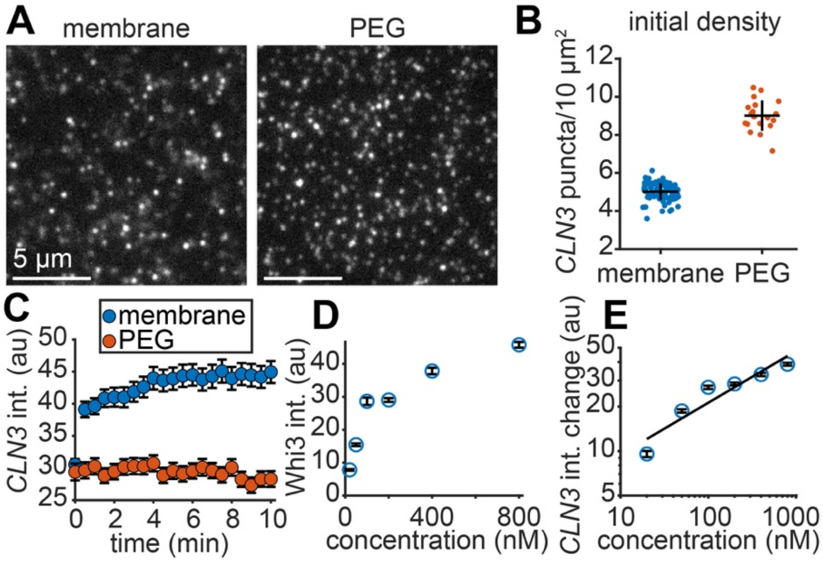


**Figure S7**. *CLN3* tethering to a diffusive membrane surface is required for Whi3 recruitment and rapid clustering into higher-order Whi3/*CLN3* assemblies. (**A**) Images of *CLN3* puncta tethered to membrane (left) or PEG surface (right) prior to addition of Whi3. Images contrasted equally. (**B**) Initial densities of *CLN3* puncta on membranes or PEG surfaces prior to addition of Whi3. (**C**) Average *CLN3* puncta intensity on membrane or PEG surfaces as a function of time after addition of 50 nM Whi3. PEG-tethered *CLN3* did not recruit Whi3 or cluster to the same extent as membrane-tethered *CLN3*, despite the higher initial *CLN3* density on PEG surfaces. (**D**) Intensity of Whi3 within membrane-associated clusters as a function of bulk Whi3 concentration. (**E**) Intensity of *CLN3* within membrane-associated clusters as a function of bulk Whi3 concentration. Data repeated from Fig. 4C, shown here on log-log axes with fit to a power law function (black line) with exponent 0.35 ± 0.18 (95% CI of fit), comparable to the power law exponent of 0.31 ± 0.08 in Fig. 4E. n = 90 and 20 images of membrane and PEG-tethered *CLN3*, respectively, in (B); n > 650 puncta per data point in (C); n > 15,000 puncta per data point in (D-E). Error bars indicate standard deviation in (B); 95% CI in (C-E). All data pooled from at least three independent experiments. SLB membrane composition: 99 mol% DOPC, 1 mol% DOPE cap-biotin.

**SI Movie Captions**

**Movie S1**. Whi3 assemblies are tethered to the ER in live *Ashbya* cells. Movie shows spinning disc confocal time-lapse of *Ashbya* hypha expressing Whi3-tdTomato (magenta) and ER marker Sec63-GFP (green). ER channel on the left, Whi3 channel in the center, and merged channels on the right, 30 s/frame.

**Movie S2**. Whi3 condensates rapidly form on a membrane in the absence of RNA. Movie shows TIRF microscopy time-lapse of planar Whi3 condensates forming on SLB following addition of 50 nM Whi3, no RNA, 30 s/frame. SLB membrane composition: 4% DGS NTA-Ni, 96% DOPC.

**Movie S3**. Whi3 condensates rapidly form on a membrane in the presence of RNA. 100 pM of Cy3-labeled *CLN3* RNA (magenta) was first added to the experiment well, followed by 50 nM of Whi3-Atto488 (green). Movie shows TIRF microscopy time-lapse of planar Whi3 condensates forming on SLB. Whi3 channel on the left, *CLN3* channel in the center, and merged channels on the right, 30 s/frame. SLB membrane composition: 4% DGS NTA-Ni, 96% DOPC.

**Movie S4**. Membrane-associated Whi3 condensates do not homogeneously incorporate RNA but instead cluster RNA at condensate edges. 100 pM of Cy3-labeled *CLN3* RNA (magenta) was first added to the experiment well, followed by 50 nM of Whi3-Atto488 (green). Movie shows TIRF microscopy time-lapse of planar Whi3 condensates forming on SLB. Whi3 channel on the left, *CLN3* channel in the center, and merged channels on the right, 30 s/frame. *CLN3* and Whi3 channels contrasted equally in Movies S4 and S5. SLB membrane composition: 4% DGS NTA-Ni, 96% DOPC.

**Movie S5**. Membrane-tethered *CLN3* RNA recruits Whi3 from solution and rapidly clusters into punctate assemblies. *CLN3* RNA (magenta) was first tethered to the membrane at a density of approximately 5 puncta/10 μm$^2$, and 50 nM Whi3 protein (green) was added in solution. Movie shows TIRF microscopy time-lapse of punctate Whi3/*CLN3* clusters forming on SLB after addition of Whi3. *CLN3* channel on the left,

Whi3 channel in the center, and merged channels on the right, 30 s/frame. *CLN3* and Whi3 channels contrasted equally in Movies S4 and S5. SLB membrane composition: 1% DOPE cap-biotin, 99% DOPC.

**Movie S6**. Immobile, PEG surface-tethered *CLN3* RNA does not recruit Whi3 to the same extent as membrane-tethered *CLN3* and does not cluster. *CLN3* RNA (magenta) was first tethered to the PEG surface at a density of approximately 9 puncta/10 $\mu m^2$, and 50 nM Whi3 protein (green) was added in solution. Movie shows TIRF microscopy time-lapse of *CLN3* puncta on PEG surface after addition of Whi3. *CLN3* channel on the left, Whi3 channel in the center, and merged channels on the right, 30 s/frame.
